# The helminth TGF-β mimic TGM4 is a modular ligand that binds CD44, CD49d and TGF-β receptors to preferentially target myeloid cells

**DOI:** 10.1101/2023.11.13.566701

**Authors:** Shashi P. Singh, Danielle J. Smyth, Kyle Cunningham, Ananya Mukundan, Chang-Hyeock Byeon, Cynthia S. Hinck, Madeleine P. J. White, Claire Ciancia, Nątalia Wosowska, Anna Sanders, Regina Jin, Sergio Lilla, Sara Zanivan, Christina Schoenherr, Gareth Inman, Maarten van Dinther, Peter ten Dijke, Andrew P. Hinck, Rick M. Maizels

## Abstract

The murine helminth parasite *Heligmosomoides polygyrus* expresses a family of modular proteins which, replicating the functional activity of the immunomodulatory cytokine TGF-β, have been named TGM (TGF-β Μimic). Multiple domains bind to different receptors, including TGF-β receptors TβRI (ALK5) and TβRII through domains 1-3, and prototypic family member TGM1 binds the cell surface co-receptor CD44 through domains 4-5. This allows TGM1 to induce T lymphocyte Foxp3 expression, characteristic of regulatory (Treg) cells, and to activate a range of TGF-β-responsive cell types. In contrast, a related protein, TGM4, targets a much more restricted cell repertoire, primarily acting on myeloid cells, with less potent effects on T cells and lacking activity on other TGF-β-responsive cell types. TGM4 binds avidly to myeloid cells by flow cytometry, and can outcompete TGM1 for cell binding. Analysis of receptor binding in comparison to TGM1 reveals a 10-fold higher affinity than TGM1 for TGFβR-I (TβRI), but a 100-fold lower affinity for TβRII through Domain 3. Consequently, TGM4 is more dependent on co-receptor binding; in addition to CD44, TGM4 also engages CD49d (Itga4) through Domains 1-3, as well as CD206 and Neuropilin-1 through Domains 4 and 5. TGM4 was found to effectively modulate macrophage populations, inhibiting lipopolysaccharide-driven inflammatory cytokine production and boosting interleukin (IL)-4-stimulated responses such as Arginase-1 *in vitro* and *in vivo*. These results reveal that the modular nature of TGMs has allowed the fine tuning of the binding affinities of the TβR- and co-receptor binding domains to establish cell specificity for TGF-β signalling in a manner that cannot be attained by the mammalian cytokine.

## Introduction

Many infectious agents exploit the pivotal host immunoregulatory pathway driven by transforming growth factor-β (TGF-β) [1,2]. In the case of helminth worm parasites, their fecundity and longevity depends upon a dampened immune system, in some cases muted by regulatory cells induced by cytokines such as TGF-β. Hence, it was remarkable to discover a helminth, *Heligmosomoides polygyrus*, that has convergently evolved a functionally active, but structurally unrelated, mimic of TGF-β (named TGM1) that binds strongly to mammalian plasma membrane TGF-β receptors [3–5]. TGM1 acts as a fully functional activator of the TGFβ signalling pathway, down-regulating inflammation in mouse models [6–9] and inducing the differentiation of both mouse and human immunosuppressive regulatory T cells (Tregs) through the canonical transcription factor Foxp3 [10,11]. The ability of this parasite to drive Treg differentiation may therefore be explained by its production of TGM1 as central part of a strategy to evade host immunity [12].

TGF-β activates cells through a heterodimeric receptor composed of two transmembrane serine/threonine kinases, TβRI (ALK5) and TβRII; its mode of binding is to first ligate TβRII, forming a complex that recruits and phosphorylates TβRI [13]. In contrast, TGM1 was found to independently bind both receptor subunits, with a particularly high affinity for TβRI [3]. TGM1 is comprised of 5 modular domains distantly related to the complement control protein (CCP) or Sushi protein family, with Domains 1 and 2 binding TβRI, while Domain 3 binds TβRII [4]. Thus, loss of any of Domains 1-3 completely ablated activity of TGM1, confirming that binding to both subunits of the receptor is required for signal transduction [14].

Recently, we ascertained that Domains 4 and 5 of TGM1 confer an additional binding specificity, for CD44, which potentiates activation of cells through the TGF-β pathway [15]. CD44 is widely expressed on hematopoietic cells, as well as some stem cell populations, interacting with extracellular matrix components such as hyaluronic acid [16]. It is prominent in immune cell types, upregulated in memory/effector T cells, [17] and includes a cytoplasmic domain capable of signal transduction. Hence, it was suggested that TGM1 has evolved to preferentially target CD44^+^ immune cells for modulation during *H. polygyrus* infection [15].

*H. polygyrus* expresses a suite of proteins related to TGM1, forming a multi-gene family of at least 10 members with up to 7 CCP-like domains [14]. Among these homologues, TGM2 and TGM3 with 93-100% identity in Domains 1-3 showed similar functional activity to TGM1 activating the signal pathway in a fibroblastic reporter cell line [14]. However, in the same assay, TGM4 was found to be inactive, despite the high sensitivity of the reporter cells [18]. and 80.2% amino acid identity to TGM1 (**Suppl. Fig. 1 A, B**).

We therefore performed a more detailed investigation of TGM4, which we now report differs from TGM1 in affinity for each TβR, and interacts with a wider range of co-receptors, resulting in a higher level of cell specificity that targets myeloid cells rather than fibroblasts. Taken together, these results highlight the importance of co-receptors, in addition to the TβRs, in delivering signals through the TGF-β pathway. This remarkable modular gene family has thereby gained the ability, unlike TGF-β itself, to selectively target different host cell populations.

## Results

### Selective activation of T cells and macrophages by TGM4

The novel TGF-β mimic (TGM) family from *H. polygyrus* was first identified by activation of the TGF-β**-**responsive fibroblast-derived reporter cell line MFB-F11, and subsequently shown to induce the transcription factor Foxp3 in naïve murine T cells [3,14]. As reported previously [3,14], TGM4 showed no activity in the same assay (**Fig. 1 a**,). However, a small but significant signal was found with a fibroblastic (NIH3T3) CAGA12 luciferase reporter cell line (**Fig. 1 b**). We therefore examined whether TGM4 could drive Foxp3 in mouse splenic T cells and found positive induction albeit at lower efficacy than TGM1 (**Fig. 1 c**), while another family member that was inactive on MFB-F11 cells (TGM7) was also negative for Foxp3 induction. Hence, TGM4 presented a uniquely discordant activity that we investigated further.

**Figure 1.**
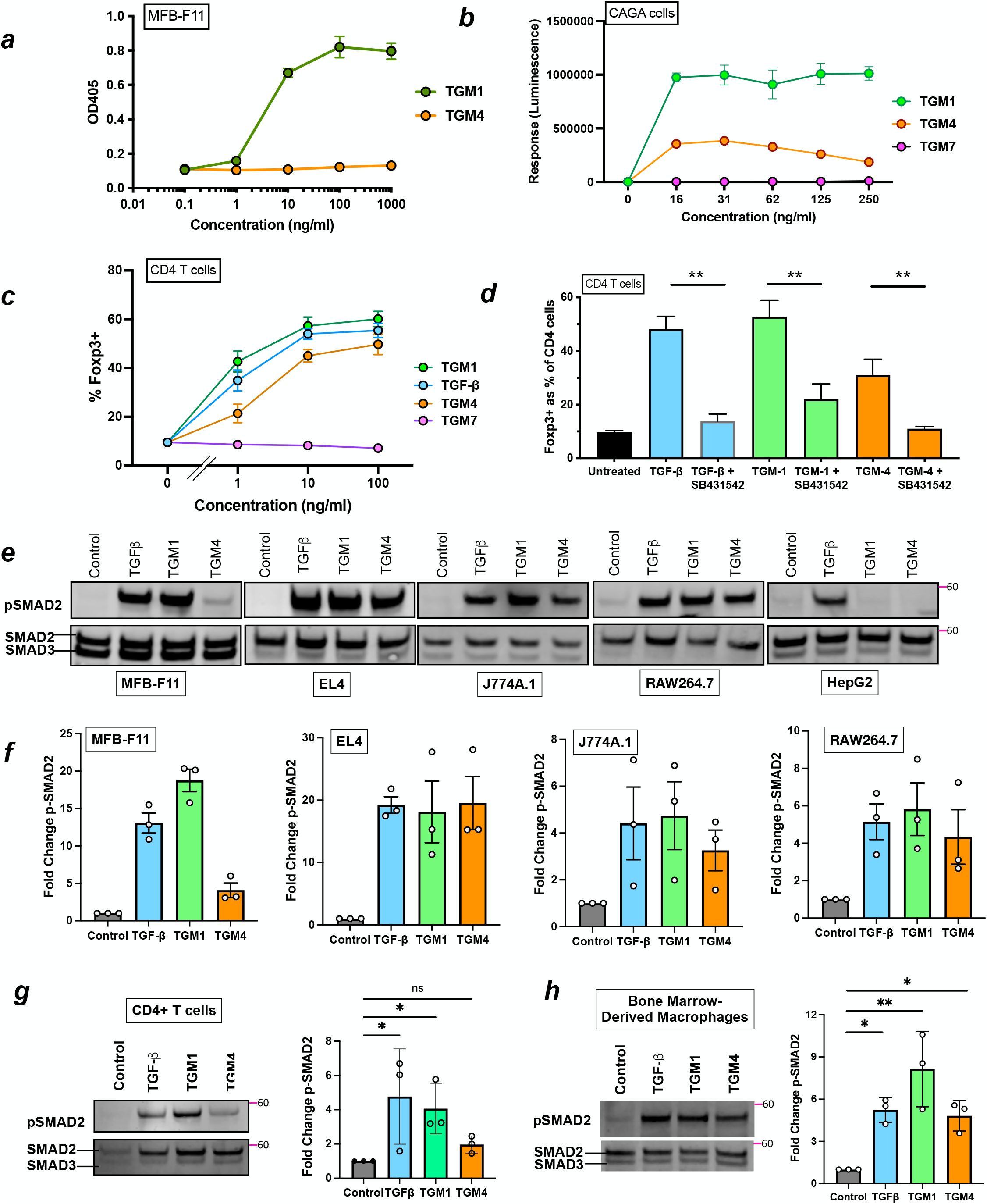
Differential activation of fibroblasts, T cells and macrophages by TGM4. ***a.*** Response of MFB-F11 reporter fibroblasts to recombinant TGM1 and TGM4 proteins, measured by an enzymatic assay (OD405 nm) for release of alkaline phosphatase. ***b.*** Response of CAGA12 luciferase reporters to TGM1 and TGM4; a more distantly related family member, TGM7, is included as a negative control). ***c.*** Induction of Foxp3 expression in mouse splenic CD4^+^ T cells incubated with TGM1, TGM4, TGM7 or TGF-β. ***d.*** Inhibition of Foxp3 induction of TGM1 and TGM4 in the presence of SB431542, which blocks kinase activity of ALK5, receptor I for TGF-β. ***e.*** SMAD2 phosphorylation in cell lines of MFB-F11 fibroblasts, EL4 T-lymphocytes, J774A.1 RAW 264.7 macrophages, and HepG2 cells stimulated with TGF-β, TGM1 and TGM4, measured by Western blotting; upper row probed with anti-pSMAD2; lower row with anti-SMAD2/3 antibody. ***f.*** Densitometric analyses of SMAD2 phosphorylation, as in ***e***, from 3 independent replicate experiments. ***g.*** SMAD2 phosphorylation in primary splenic CD4^+^ T cells, assessed as in ***e*** and ***f***, by Western blot (left panel) and by densitometric analyses if 3 independent experiments (right panel). ***h.*** SMAD2 phosphorylation in bone marrow-derived macrophages, as in ***g***. * p<0.05, ** p<0.01.

To confirm that TGM4 was, like TGM1, acting through the TGF-β signalling pathway in T cells, cultures were supplemented with the small molecule inhibitor SB431542 which blocks the kinase activity of ALK-4, 5 (TαRI) and 7 [19]. As shown in **Fig. 1 d**, while TGM4 was less potent than TGM1 in inducing Foxp3, both ligands were fully suppressed by addition of SB431542, as previously established for TGM1 [3].

We then probed different cell types for TGM1- and TGM4-stimulated phosphorylation of SMAD2 that is immediately downstream of the TβRI ALK5 kinase in the signalling pathway. Cell lines were incubated with these ligands, and mammalian TGF-β, for 60 minutes, then lysed and analysed by Western blot with anti-SMAD2 and anti-phospho-SMAD2 (p-SMAD2) antibodies. As shown in **Fig 1e, f**, TGM-4-stimulated MFB-F11 fibroblasts showed only low levels of SMAD2 phosphorylation in contrast to TGM1 and TGF-β. However, in the EL4 T cell line, and in macrophage lines (J774A.1 and RAW264.7) all three ligands were equally active. We also tested a hepatoma cell line, HepG2, which responded only to TGF-β (**Fig 1 e**).

We then investigated responses of primary murine hematopoietic cells, measuring SMAD2 phosphorylation 1 hr following stimulation with the different ligands. When CD4^+^ murine splenic T cells were stimulated with TGM4, responses were much weaker than with TGF-β or TGM-1, and indeed did not attain statistical significance (**Fig. 1 g**). As similar cells were capable of Foxp3 induction after 72 hrs co-incubation with TGM4 (**Fig. 1 c**), the possibility was raised that activation follows a slower time course, as indeed observed for TGM1 compared to TGF-β [11]. To test this, we employed imaging flow cytometry to measure nuclear localisation of SMAD2/3 in splenic T cells at 1 and 16 hrs post-stimulation; although at the earlier time point TGM4-stimulated T cells were at baseline values (**Suppl. Fig. 2 a**), by 16 hrs they were elevated and comparable to cells activated with the other ligands (**Suppl. Fig. 2 b**). Thus, while TGM4 has a relatively subdued ability to activate TGF-β signalling in primary T cells, it is sufficient to induce measurable responses over a 16-72 hr time frame.

We similarly analyzed SMAD activation in bone marrow-derived macrophages; in this subset, responses to each ligand were comparable, with TGM4 also inducing a significant level of pSMAD2/3 measured by Western blot (**Fig. 1 h**) and SMAD2/3 nuclear localisation (**Suppl. Fig. 2 c**) within 1 hour of stimulation. In additional analyses, TGM4 was found to lack activity on other epithelial (NM18) and fibroblast (NIH 3T3) cell lines (**Suppl. Fig. 2d, e**), but did induce SMAD2 phosphorylation in dendritic cells, represented both as cell lines (MuTu, D1) and bone marrow-derived DCs (**Suppl. Fig. 2 g, h**). Hence, TGM4 displays a strong predilection for cells of the myeloid lineage.

### TGF-β Receptor Binding by TGM4

We next tested whether activation of the SMAD signalling pathway by TGM4 could be induced by the same domains (D1/2/3) that are required for activation by TGM1 [14]. Cells incubated with full-length (D1/2/3/4/5) or truncated (D1-3 or D4-5) portions of TGM1 and TGM4 were probed for SMAD2 phosphorylation. As shown in **Fig. 2 a**, only full-length or D1-3 of TGM1 elicited a p-SMAD response in MFB-F11 fibroblasts, but neither full-length nor D1-3 of TGM4 did so. In contrast, when a macrophage cell line, RAW 264.7, was tested, both proteins drove p-SMAD, and for both ligands activity resided, albeit attenuated, in D1-3 (**Fig. 2 b**).

**Figure 2.**
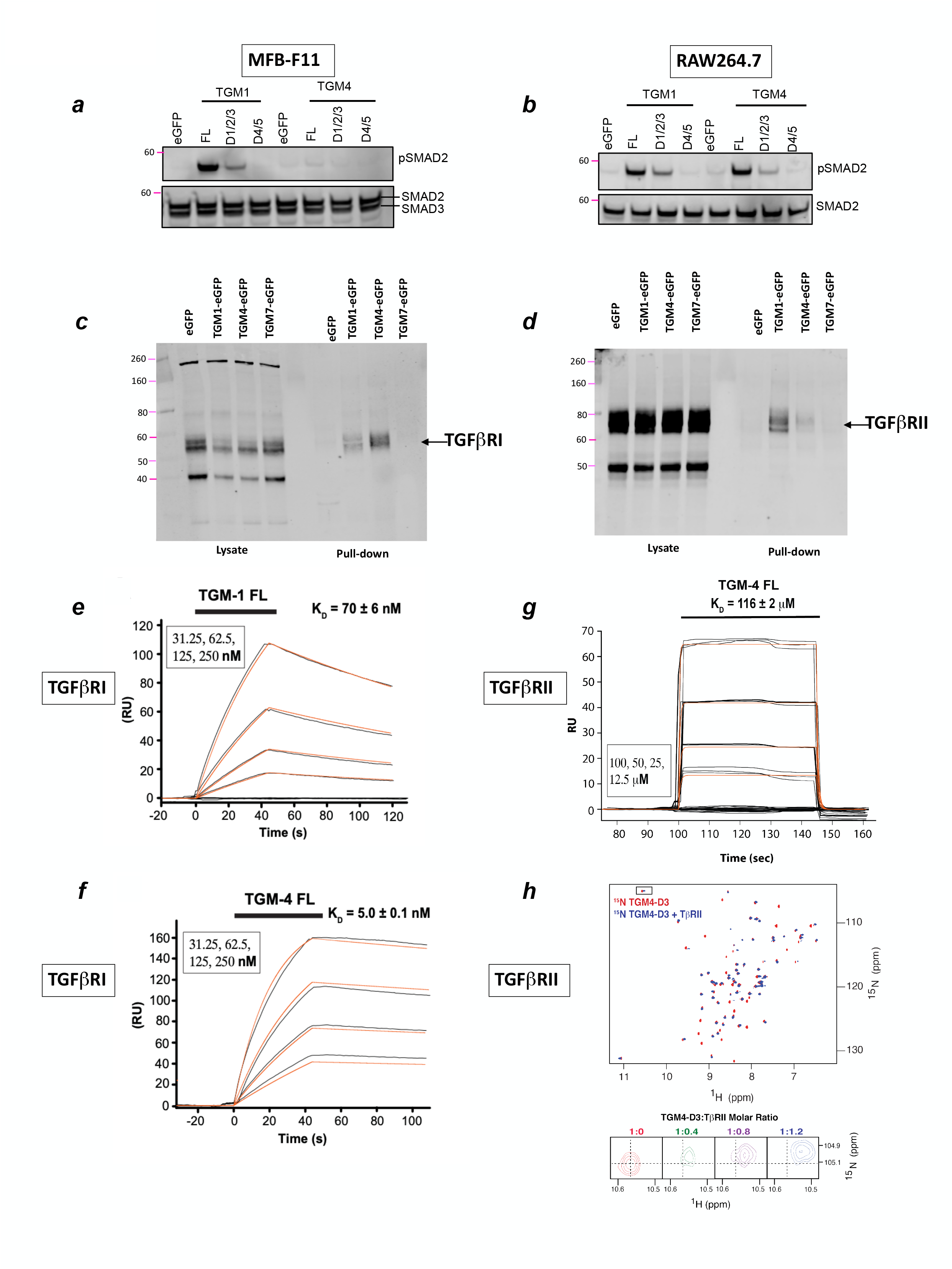
TGM4 Domain and Receptor Binding Analysis. ***a, b***. SMAD2 phosphorylation induced by Domains1/2/3 by both TGM1 and TGM4 when added to MFB-F11 fibroblasts (***a***) or RAW264.7 macrophage cell line (***b***). Serum starved cells were incubated with conditioned medium from eGFP alone, TGM4 D1-5, D1-3 or D4-5 transfected cells for 1 h and analysed for pSMAD2 and SMAD2/3 by Western blotting. ***c, d.*** TβRI (***c***, TGFβRI) and TβRII (***d***, TGFβRII) GFP-TRAP pull down from MFB-F11 cells transfected with eGFP alone, or eGFP fused to TGM1, TGM4 or TGM7; whole cell lysates shown in left-hand side of each panel, and anti-GFP immunoprecipitates (pull downs) shown in right hand side. Western blots with anti-TβRI (***c***) and anti-TGβRII (***d***) are presented. ***e,f.*** Surface plasmon resonance (SPR) sensorgrams of full-length TGM1 and TGM4 binding to biotinylated Avi-tagged TβRI immobilized on a streptavidin chip. Injections were performed as a two-fold dilution series and are shown in black, with the orange traces over the raw data showing curves fitted to a 1:1 model, when possible. The black bars over the top of the sensorgrams correlates to the injection period, and the injection concentrations are on the top left of each sensorgram. ***g.*** SPR sensorgram of full-length TGM4 binding to biotinylated Avi-tagged TβRII immobilized on a streptavidin chip, with injection performed as above; injection period depicted by the black bar at top, and injection concentrations at bottom left. ***h.*** Nuclear magnetic resonance analysis of TGM4 Domain 3 binding to TβRII; D3 was labelled with ^15^N at a concentration of 100 µM in buffer consisting of 25 mM sodium phosphate, 50 mM sodium chloride, 5% ^2^H2O, pH 6.0. ^1^H-^15^N spectra of ^15^N-TGM4 D3 (red) overlaid with the ^1^H-^15^N spectra of the same protein bound to 1.2 molar equivalents of unlabeled TβRII (blue) at 37°C. Expansion of intermediate titration points (1:0, 1:0.4, 1:0.8, 1:1.2 ^15^N-TGM4 D3:TβRII) of the boxed residues are shown in the lower panel. Data showing no interactions with other Type II receptors are presented in **Suppl. Fig. 3**.

To investigate whether TGM4 differs from TGM1 in its interactions with the two TGF-β receptor chains (TβRI and TβRII), we used a system of endogenous expression of enhanced green fluorescent protein (eGFP)-TGM fusion proteins in cell lines, followed by anti-eGFP antibody pull-down of the ligand and any associated receptors. In this manner, we identified that TGM1 and TGM4, but not TGM7, forms complexes with both TβRI (**Fig. 2 c**) and TβRII (**Fig. 2 d**). While TGM4 co-precipitated TβRI more strongly than did TGM1, the interaction with TβRII was substantially weaker than observed with TGM1.

To ascertain whether TGM4 binds directly to each receptor chain, and to understand the respective receptor affinities of TGM1 and TGM4, surface plasmon resonance (SPR) measurements were made for binding to TβRI and TβRII. SPR analysis showed substantially higher affinity of TGM4 than TGM1 (**Fig. 2 e, f**) for TβRI, estimated at 3-5 nM vs 70 nM respectively (**Table 1, Suppl. Table 1**). Notably, TGM4 binding to TβRI shows a faster on-rate and slower off-rate than does TGM1. Unlike the direct binding of TGMs to TβRI, it is well established that TGF-β has little direct affinity for TβRI unless complexed to TβRII [20].

**Table 1.** Summary of binding affinities of TGM1 and TGM4 for TβRI, TβRII, and CD44.

| TGM Ligand | Measurement | TβRI | TβRII | mCD44 | hCD44 |
| --- | --- | --- | --- | --- | --- |
| TGM1-FL | SPR | 70 ± 6 nM | 0.61 ± 0.01 mM<sup>1</sup> | | |
| TGM4-FL | SPR | 3.2 ± 0.1 nM | 116 ± 2 μM | | |
| TGM1-D45 | ITC |  |  | 62 (28, 117) nM<sup>2</sup> | 140 (86, 221) nM<sup>2</sup> |
| TGM4-D45 | ITC |  |  | 20 (7, 41) nM<sup>2</sup> | 56 (28, 99) nM<sup>2</sup> |
| <sup>1</sup>Value previously reported by Mukundan *et al* [4].<br><sup>2</sup>Values in parenthesis represent estimated upper and lower bounds estimated by the Sedphat fitting algorithm |  |  |  |  |  |

In contrast to the higher affinity of TGM4 for TβRI, binding to TβRII was found to be weak by SPR, at 116 µM, representing >100-fold lower affinity than TGM1 (**Fig. 2 g, Table 1, Suppl. Table 1**). NMR analysis showed small but significant shifts of several signals when^15^N-labeled Domain 3 of TGM4 was titrated with increasing concentrations of TβRII (**Fig. 2 h**), also indicative of no more than moderate binding affinity. As this raised the possibility that the physiological target of TGM4 is another type II receptor, we performed SPR assays with ActRII and BMPRII, two major receptors of this type. However, we found no evidence of direct binding to either of these receptors (**Suppl. Fig. 3, a, b**). In addition, no interactions were observed between Domain 2 and the type I Activin receptor, ALK4 (**Suppl. Fig. 3 c**). We therefore concluded that the cognate receptors for TGM4 are TβRI (ALK5) and TβRII, similar to TGM1, although the two parasite proteins differ markedly, and reciprocally, in affinity for these receptors.

### CD44 Binding by TGM4

TGM1 and -4 share the same 5-domain structure (**Suppl. Fig. 1 a**); in the case of TGM1 we showed by truncation analysis that only Domains 1-3 were essential for biological activity [14]; however, Domains 4 and 5 (D4/5) enhance the potency of TGM1, binding to a cell surface co-receptor identified as CD44 [15]. To determine if TGM4 also binds CD44, we expressed eGFP-tagged TGM1 and TGM4 in MFB-F11, RAW264.7 and HepG2 cells, followed by GFP-TRAP pulldown and Western blotting analyses. As shown in **Figure 3 a**, a strong CD44 band was observed with both TGM1 and TGM4 in the fibroblast and macrophage cell lines, but not in hepatocytes. Quantification from replicate Western blot experiments showed that CD44 was considerably more prominent in RAW264.7 cells than MFB-F11 (**Figure 3 b**). In addition, higher TβRI levels were co-precipitated with TGM4 than TGM1 in RAW264.7 cells (**Figure 3 a, c**). In contrast, TβRII co-precipitation was significantly greater with TGM1 (**Figure 3 a, d**), consistent with the weak TGM4 binding to this receptor noted above. Taken together, these data suggest that the strength of interaction with CD44 and both TβRs could be related to the differential activity of TGM4 on these two cell types.

**Figure 3.**
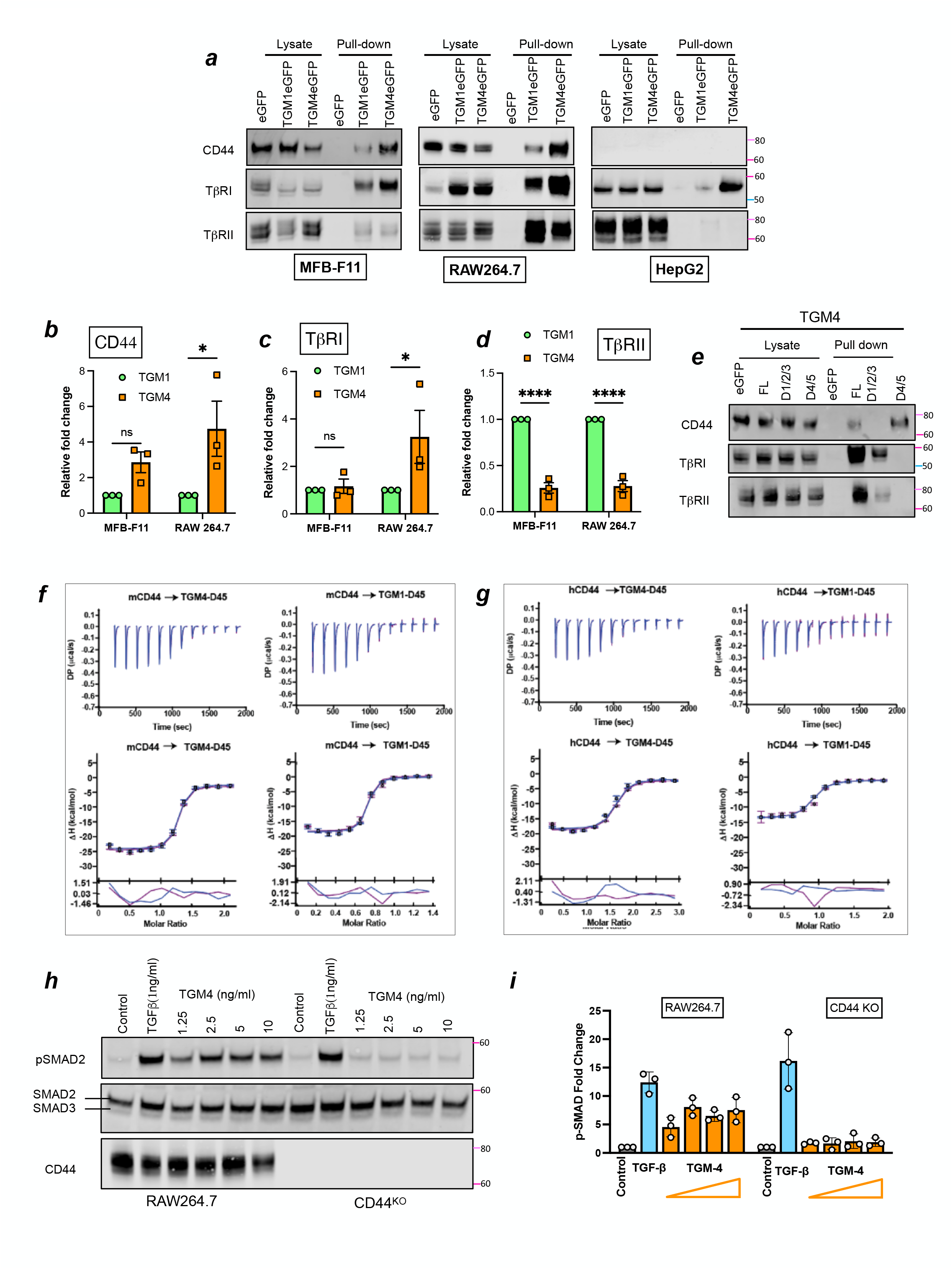
TGM4, like TGM1, binds CD44. ***a.*** eGFP trap pull down and Western blotting analyses of MFB-F11, RAW264.7 and HepG2 cells transfected with eGFP alone, or with eGFP-TGM1 or TGM4 fusions; whole cell lysates shown in the left-hand side of each panel, and anti-GFP immunoprecipitates (pull downs) shown on the right; Western blots were probed with antibodies to CD44, TβRI, and TβRII ***b-d.*** Densitometric analysis of pulldown proteins CD44 (**b**), TβRI (**c)**, and TβRII (**d**) from MFB-F11 or RAW264.7 cells expressing TGM1 or TGM4. * p<0.05, *** p<0.001. ***e.*** eGFP trap, pulldown and Western blotting (in what cells?) as in ***a***, with TGM4 full-length (FL) and truncated constructs D1/2/3 and D4/5. ***f,g***. ITC binding isotherms of TGM4 D4/5 binding to murine and human CD44 (**f** and **g**, respectively). In comparison, similar analyses of TGM1 D4/5 binding are also shown on the right-hand side of each panel. ***h,i***. TGM4 activation of pSMAD signalling in RAW264.6 macrophages is dependent on CD44 expression, shown as exemplar Western blot (***h***), and densitometric data from 3 independent experiments (***i***); in (***i***) TGF-β is shown to activate in a CD44-independent manner.

To dissect interactions of TGM4 with each receptor at the domain level, eGFP fusions of truncated proteins were designed, expressing D1-3 or D4-5 of TGM1 and TGM4 in RAW264.7 cells. Lysates of cells expressing each construct were immunoprecipitated using GFP-TRAP beads and analysed by Western blotting. D1-3 constructs precipitated the TβRs, but not CD44, while when D4-5 was expressed, the converse was true (**Fig. 3 e**), as recently reported for TGM1 [15].

To more precisely evaluate TGM4-CD44 interactions, the binding of TGM1 and TGM4 with recombinant human and mouse CD44 was measured using isothermal titration calorimetry (ITC). For these measurements, the D4-5 fragments of each TGM were employed, as this segment of TGM1 carries the CD44 binding capacity [15]. As shown in **Figure 3 f, g**, both helminth proteins bound each CD44 molecule with similar affinities, which were determined to be in the 50-200 nM range (**Table 1, Suppl. Table 2**).

To establish whether CD44 was essential for TGM4 activation of myeloid cells, we tested control RAW264.7 macrophages, and a line derived from these cells from which CD44 had been genetically deleted by CRISPR-Cas9. As shown in **Fig. 3 h, i**, SMAD phosphorylation was reduced to background levels in macrophages lacking CD44, when tested with a range of concentrations of TGM4 that elicited strong responses in CD44-sufficient cells.

### Preferential Binding of TGM4 to Myeloid Cells

We recently reported that Alexa Fluor-488 (AF488)-labelled TGM1 binds strongly to the surface of MFB-F11 and EL4 cells, as measured by flow cytometry [15]. We noted that TGM4 bound these, and other, cell lines with greater intensity than seen with TGM1 (**Suppl Fig. 4a**). Preferential binding of TGM4 was most evident on two macrophage cell lines (J774A-1 and RAW264.7), while no binding was observed to the hepatocyte line, HepG2. Notably, the intensity of binding to J774 macrophages was significantly higher than each of the other immune-derived lines.

We then tested TGM1 and TGM4 binding to primary peritoneal exudate cells, by flow cytometry using AF594-labelled proteins. Both ligands showed extensive binding to CD3^+^ T cells and CD11b^+^F4/80^+^MHC-II^low/-^ tissue resident macrophages (**Fig. 4 a**), and in each case staining was more intense by TGM4 compared to TGM1 (**Fig. 4 b**). Staining of T lymphocytes was quite heterogenous, while 100% of macrophages were bound by TGM4.

**Figure 4.**
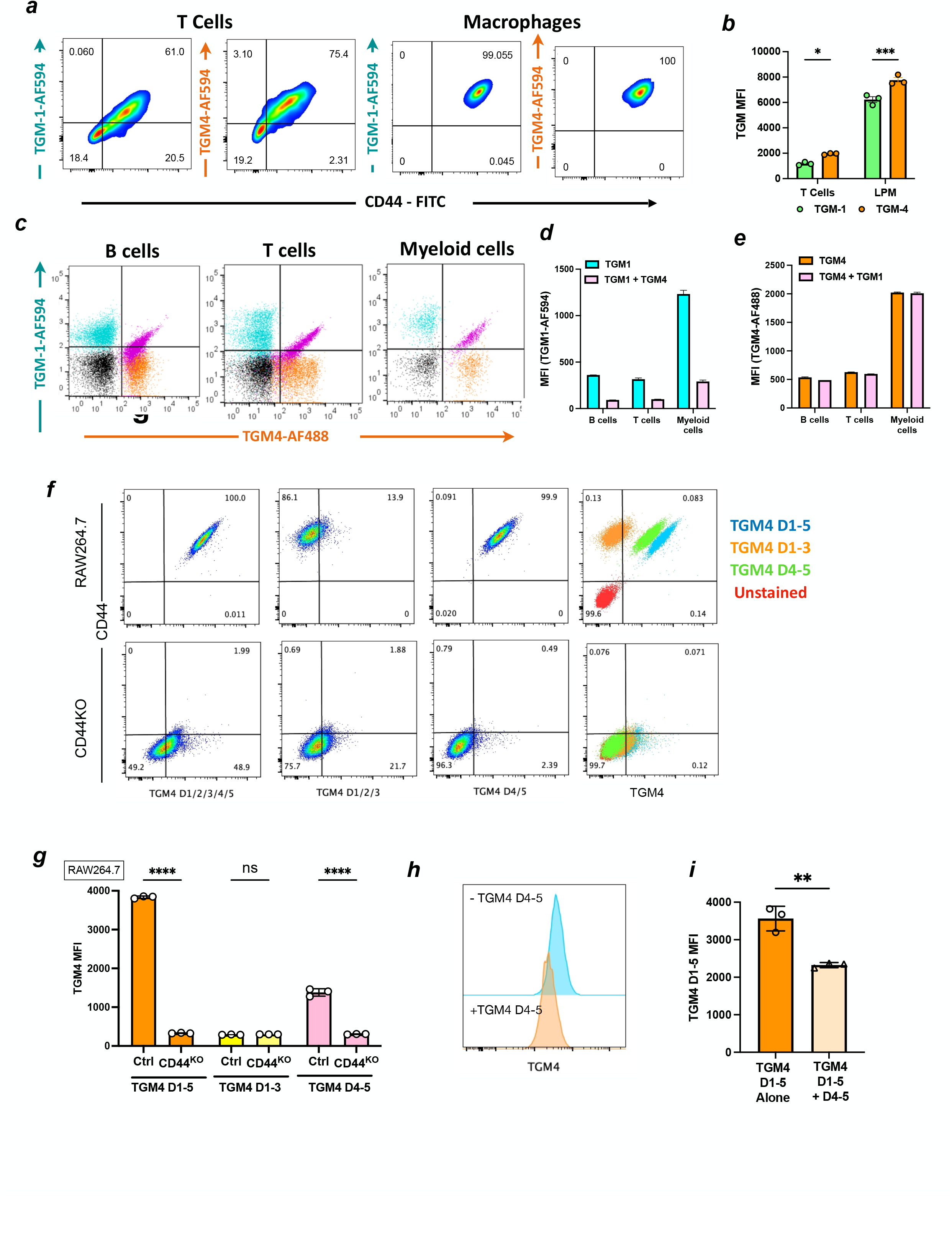
TGM4 binding to host immune cells. ***a.*** Co-staining of CD44 and Alexa Fluor 594 (AF594)-labelled TGM1 or TGM4 to peritoneal CD3^+^ T cells, and F4/80+ large peritoneal macrophages, measured by flow cytometry. Percentages of the target populations in each quadrant are shown. ***b.*** Mean fluorescent intensity (MFI) for AF594 labelled TGM1 or TGM4 on replicate populations of peritoneal T cells and large peritoneal macrophages. *, p<0.05; *** p<0.001. ***c.*** Co-staining of Alexa-Fluor 594 (AF594)-labelled TGM1 and AF488-labelled TGM4 to spleen cell populations. Plots are superimposed from the indicated cell populations stained with TGM1 (cyan), TGM4 (orange), both TGM1 and TGM4 (magenta) or unstained control (black). ***d,e*** Quantification of staining by TGM1 (***d***) or TGM4 (***e***) in the absence of presence of TGM4 or TGM1 respectively, as measured by mean fluorescent intensity (MFI). ***f.*** Flow cytometric analysis of TGM4 binding to RAW264.7 wild-type and CD44-deficient cells, probed with full length TGM4 D1-5, and truncated constructs D1-3 and D4-5. Right hand panels show the 3 datasets superimposed, with full-length D1-5 (blue), D1- 3 (orange), D4-5(green), together with unstained control (red). ***g.*** Mean fluorescent intensity (MFI) in 3 independent replicate experiments comparing binding of the TGM4 full-length and truncation constructs to wild-type and CD44- deficient RAW264.7cells. ****, p<0.0001. ***h.*** Partial inhibition of AF AF488 FL TGM4 binding to RAW264.7 in presence of unlabelled TGM4 D4-5 shown as exemplar histogram (***e***) and data from 3 replicate samples (***f***). **, p<0.01.

Although TGM4 showed more intense staining, it is possible that differences in efficiency of labelling or protein stability affected by coupling Alexa-Fluor to exposed lysine residues could be responsible. However, when we co-stained with both proteins (**Fig. 4 c**), TGM4 was able to outcompete TGM1, reducing the TGM1 signal (**Fig. 4 d**) while the TGM4 signal was unaffected by the presence of TGM1 **(Fig. 4 e)**. Although weaker, TGM1 binding correlated closely with TGM4 (giving a diagonal profile in doubly-stained cells) indicating that the two ligands bind similar populations of host cells. Taken together, these data demonstrate that TGM4 is more avid than TGM1 in surface binding to immune system cells, with a strong affinity for myeloid cells.

To examine the role of CD44 in surface interactions, we deleted the *Cd44* gene in RAW264.7 cells by CRISPR-Cas9 (**Suppl. Fig 4 b**). We then tested CD44-sufficient and-deficient cells by flow cytometry with AF488-labelled full-length and truncated TGM4 proteins comprising D1-3 or D4-5. As shown in **Fig. 4 f**, we found that in the absence of either CD44, or D4-5, cell surface binding is effectively abolished. Thus, on CD44-sufficient RAW264.7 cells (upper row of **Fig. 4 f**), binding of full-length TGM4, or of D4-5, correlates closely with expression of CD44, while D1-3 shows only a low level of fluorescence; in the absence of CD44 expression (lower row) a similar residual level of binding is observed in full-length TGM4 and D1-3, suggesting this represents interactions with the TβRI/II proteins. It was also noted that the intensity of binding (MFI) of D4-5 was attenuated compared to full-length TGM4 (**Fig. 4 f**, top right panel; **Fig. 4 g**). Similar data were obtained with MFB-F11 cells with intact or deleted CD44 expression (**Suppl. Fig 4 c, d**). As with RAW264.7 cells, D4/5 binding to MFB-F11 was lower than full-length TGM4, suggesting that D1-3 may contribute to the overall binding affinity for CD44. A similar involvement of D1-3 in optimal CD44 binding was also indicated by the ability of the unlabelled D4-5 to diminish but not fully inhibit the binding of full-length AF488-labelled TGM4 to RAW264.7 cells (**Fig. 4 h, i**).

In the case of TGM1, Domain 1 was found to contribute to the ability of domain 2 to bind TβRI, [4], explaining earlier data that Domains 1-3 were all essential for biological activity in the MFB-F11 assay [14]. A similar truncation analysis for TGM-4 was performed, evaluating pSMAD2/3 activation, indicating a parallel dependency, as in the absence of any of the first 3 domains, signalling was ablated, while in the absence of D4 and/or D5, an attenuated level of signalling could be detected (**Suppl. Fig. 4 e**).

### TGM4 Binds Additional Co-Receptors

To better understand why TGM4 binds cells more avidly than TGM1 despite a similar affinity for CD44 within D4-5, we investigated the possibility that TGM4 interacts with additional co-receptors not recognized by TGM1. Pull-down of TGM4-binding proteins from splenocytes revealed, in addition to CD44 and TGM4 itself, 4 more candidates: integrin α4 (CD49d), CD72, Mrc1 (CD206) and Lirb3 (**Fig. 5 a**). While CD44 was evident in pull-downs from MFB-F11 cells, none of these other candidates were apparent (**Fig. 5 b**). We also performed similar procedures on the macrophage cell line J774, confirming that TGM4 interacted with CD49d and CD72, in addition to CD44 (**Fig. 5 c**). In addition, in all cells probed with TGM4, neuropilin- 1 (Nrp-1) was identified. Notably, parallel analyses of the same three cell populations with TGM1 failed to show interactions with these CD49d, CD72 or Nrp-1 (**Fig. 5 d-f**).

**Figure 5.**
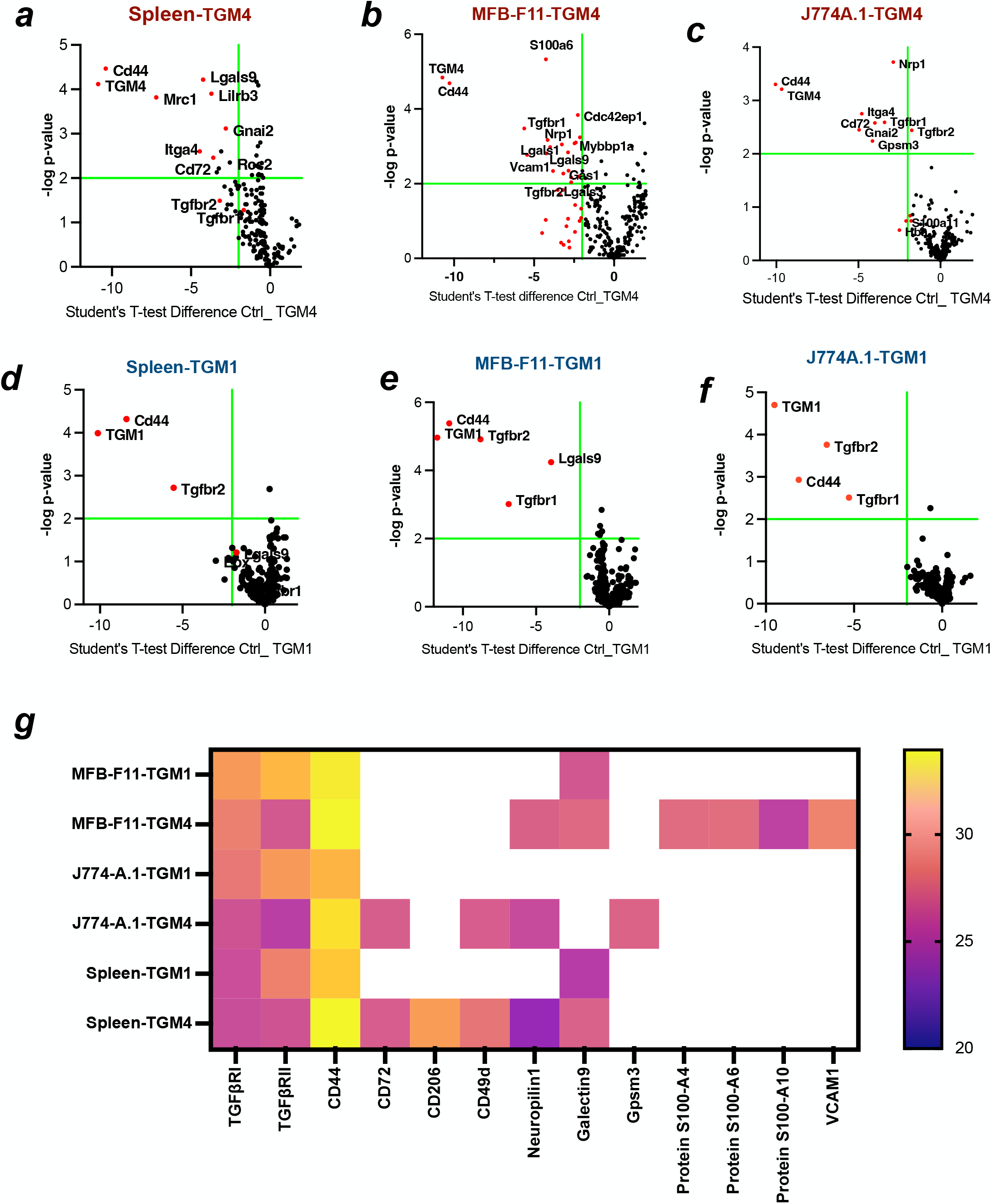
Pull down and mass spectrometric identification of novel co-receptors for TGM4. ***a-c.*** TGM4 pull down samples were subjected to mass spectrometry and analysed relative to control samples, in 3 independent experiments), using C57BL/6 strain murine splenocytes (***a***), MFB-F11 cells (***b***) and J774 macrophages (***c***). ***d-f*** Parallel analyses of pull down samples in the same experiments with TGM1 in the 3 indicated cell types. ***g.*** Heat map of proteins detected in each case.

Taken together, these data demonstrated that TGM4 associates not only with the CD44 co-receptor, but also CD49d and CD72 which were not detected on fibroblasts, and with Nrp-1 which was expressed in all cell types studied (**Fig. 5 g**).

#### CD44-Dependent and Independent Co-Receptors

In co-precipitation experiments, partner proteins may interact with different members of a complex. To ascertain whether TGMs directly bind co-receptor proteins, or do so in conjunction with CD44, we performed pulldown experiments in RAW264.7 cells with unaltered or deleted *Cd44*. In CD44-sufficient cells, CD49d, CD206 and Nrp-1 were all detectable in precipitates from cells incubated with biotinylated TGM4, but not TGM1 (**Fig. 6 a**); the loss of CD44 from RAW264.7 cells ablated CD206 and Nrp-1 detection, but CD49d remained present in the TGM4 pull down. Hence although CD206 and Nrp-1 are found only in TGM4 complexes, their presence is dependent on CD44. A parallel result was observed in MFB-F11 cells, in which CD49d is not expressed, but NRP1 is present; as with RAW264.7 cells, NRP1 is precipitated only in CD44-sufficient cells (**Fig. 6 b**).

**Figure 6.**
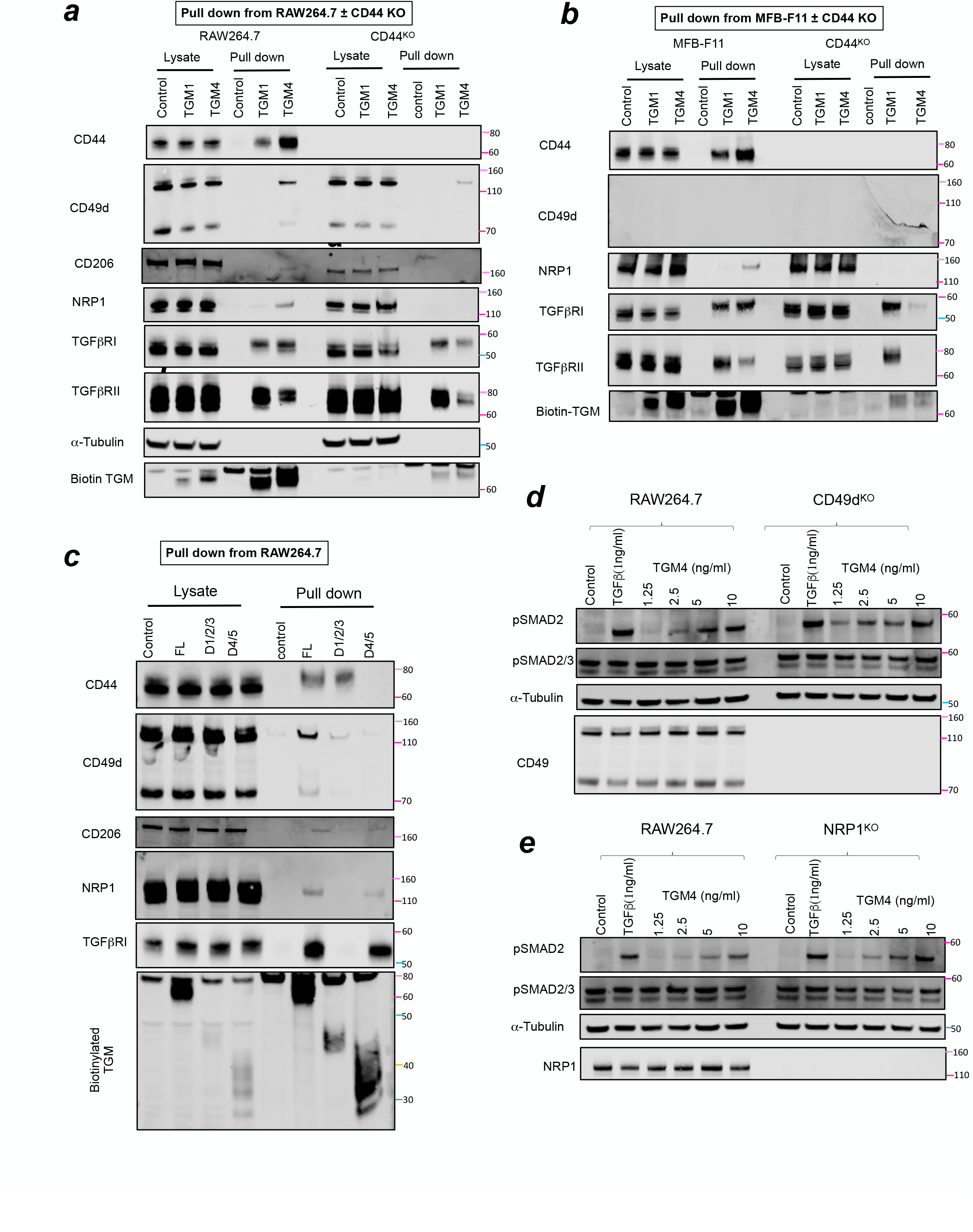
Validation of CD49d, CD72, CD206 and Nrp-1 as co-receptors for TGM4. ***a.*** Pull down and Western blot analysis in CD44-sufficient and deficient RAW264.7 cells, probed with antibodies to the indicated proteins. ***b.*** As ***a***, but with MFB-F11 cells sufficient or deficient for CD44. ***c.*** As ***a***, but comparing full-length and truncated constructs of CD44 by pulldown and Western with antibodies to indicated proteins. ***d.*** SMAD phosphorylation in RAW264.7 cells sufficient or deficient for CD49d, following stimulation with the indicated concentrations ofr TGM4. e. As ***d***, but with RAW264.7 cells sufficient or deficient for NRP1.

We next asked whether the CD44-binding domains D4-5 are required for association with these three co-receptors; the same streptavidin pull down system was used with cells incubated with biotin labelled full-length (D1-5), D1-3 or D4-5 TGM4. CD206 and NRP1 were found only in full-length and D4-5 constructs, consistent with them being dependent on CD44. In contrast, CD49d was associated with D1-3 proteins, which also bind the two TGF-β receptors, albeit the binding is weaker than with full-length TGM4 (**Fig. 6 c**).

CD49d and Nrp-1 knockout cells were then tested to determine if either gene was required for signalling in response to TGM4. These cells, and unmodified controls, were stimulated with TGM4 for 30 minutes, and cell lysates probed for SMAD phosphorylation by Western blot. However, in neither case did gene deletion reduce responses measured by pSMAD (**Fig. 6 d,e**). Moreover, when KO cells were tested by flow cytometry, no diminution of binding by TGM4 was noted (**Suppl. Fig 5**).

### TGM4 Efficiently Modulates Macrophage Function

To test whether the preferential targeting of myeloid cells by TGM4 has functional consequences, we first examined murine macrophage cell lines, following in vitro stimulation with lipopolysaccharide (LPS). As shown in **Fig 7 a,** the inflammatory response of RAW264.7 cells as measured by interleukin (IL)-1β secretion, was inhibited to an equal degree by TGM1, TGM4 and TGF-β, while in the same cell cultures, all ligands enhanced IL-10 production (**Fig. 7 b**).

**Figure 7.**
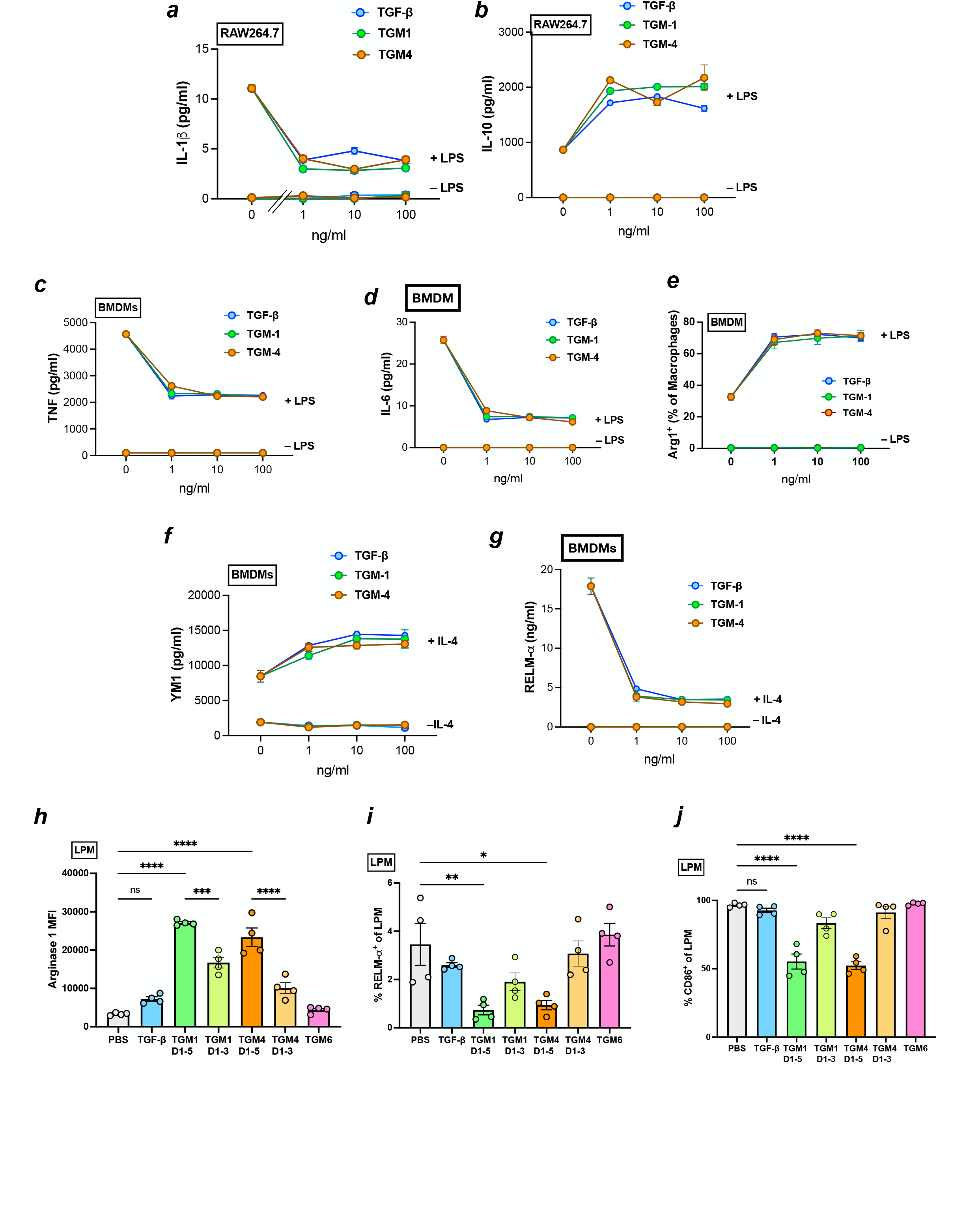
TGM4 activity on macrophage populations. ***a,b***. *In vitro* responses of RAW264.7 cells to a range of concentrations of TGF-β, TGM1 and TGM4, in the presence or absence of 100 ng/ml LPS, measured by release of IL- 1β (***a***), and IL-10 (**b**) after 24 hours of culture. ***c,d.*** In vitro responses of bone marrow-derived macrophages to a range of concentrations of TGF-β, TGM1 and TGM4, in the presence or absence of 100 ng/ml LPS, measured by release of TNF (***d***) and IL-6 (***e***) after 24 hours of culture. ***e, f,g***. Responses of bone marrow-derived macrophages to a range of concentrations of TGF-β, TGM1 and TGM4, in the presence or absence of 20 ng/ml IL-4, measured by release of Arginase-1 (***f***), Chi3L3 (Ym1) (***g***), and RELM-α (**h**) after 24 hours of culture. ***h, i***. Phenotype of resident large peritoneal macrophages, collected from the peritoneal cavity 24 hours after i.p. injection of PBS, TGF-β, TGM1 or TGM4, and analysed by flow cytometry for staining for Arginase-1 (***i***), RELM-α (***h***), and CD86 (***k***).

We then tested bone marrow-derived macrophages (BMDM) in a similar setting. TGM4 effectively inhibited the LPS-induced scretion of both TNF (**Fig 7 c)**, and IL-6 (**Fig 7 d**). Furthermore, the macrophage response to IL-4 is characterized by induction of M2 genes such as Arginase-1 [21]., chitinase-like protein Chi3L3 (Ym1) and resistin-like molecule α (RELMα) [22,23]. When TGF-β or TGMs are co-administered with IL-4, however, Arginase-1 (**Fig 7 e**) and Chi3L3 (**Fig 7 f**) expression are each markedly amplified while RELMα secretion is inhibited (**Fig 7 g**), in an uncoupling of these markers that are often considered co-ordinately produced.

Finally, to question whether TGM4 would have similar effects on primary macrophages *in vivo*, we administered recombinant proteins into the peritoneal cavity of BALB/c mice and recovered myeloid populations 24 hrs later. We focused on the resident F4/80 ^hi,^ MHCII ^lo^ peritoneal macrophage population, using flow cytometry, co-staining for type 2 macrophage markers, revealing a significant increase in Arginase-1 expression (**Fig 7 h**), and repression of RELMα (**Fig. 7 i**) that had been observed in BMDMs *in vitro,* and in each case the effects were attentuated in the absence of D4-5/CD44 interactions. Further examination of surface markers revealed a significant reduction in CD86 expression by TGM4 within the large peritoneal (resident) macrophage LPM subset (**Fig 7 j**), again contingent upon the presence of the co-receptor binding domains 4 and 5.

## Discussion

Pathogens have dramatically shaped the evolution of the immune system, but in the case of TGM protein family it appears that host immunity has driven a novel elaboration to confer cell-selectivity for a set of cytokine mimics. In mimicking TGF-β, *H. polygyrus* may have adopted a successful strategy for immune evasion that is shared, in different fashion, by many diverse infectious agents [2]. The emergence of multiple homologues which form their own multigene family may represent redundancy (for example, if the host generates antibodies to one form, the others are still effective) or specialization (such that different family members fulfill different functions and/or target different host tissues). The latter may hold true in some cases at least, as 4 of the family members (TGM7 to 10) are only expressed by larval stages in the intestinal submucosa, and not by luminal-dwelling adults [14].

The TGM family are modular constructs with between 3 and 7 homology domains all distantly related to the CCP or Sushi family (Pfam00084), modified by short insertions that evidently allow evolutionary flexibility and the ability to bind novel interaction partners [4]. Both TGM1 and TGM4 are 5-domain proteins and we find here that they share multi-specificity mediated through separate domains, Domain 2 binding TβRI and Domain 3 binding TβRII. Although the corresponding domains of the two proteins are highly conserved (86-90% amino acid identity), we find sharply contrasting binding affinities, with TGM4 being tenfold stronger for TβRI, yet 100-fold weaker in binding TβRII.

These differences may account for the inability of TGM4 to activate a fibroblast reporter line, while retaining some ability to induce T cell expression of Foxp3, the canonical transcription factor that specifies suppressive Tregs which are expanded during *H. polygyrus* infection [24,25]. However, TGM4 is markedly more active on myeloid cells than other cell types; this is perhaps not surprising given the pivotal role of macrophages and neutrophils in priming and mediating protective immunity to *H. polygyrus* [26–30]. In future studies, it would be interesting to target TGM4 by active or passive immunisation and monitor the effects on myeloid cell subsets during parasite infection.

An intriguing question is why TGM4 has evolved a lower affinity for TβRII than other members of the gene family. Οne answer may be that this, together with the recruitment of co-receptors, allows the ligand to be more discriminatory. It is likely that the selectivity of TGM4 is based on a continuum threshold; although MFB-F11 fibroblasts do express the CD44 co-receptor (and can bind TGM4 in flow cytometry assays), either the expression level is too low, and/or the additional coreceptors required for high affinity interactions are absent, resulting in the failure of TGM4 to assemble an activation complex. The ability of TGM1 to stimulate fibroblasts can thus be attributed to its >10-fold higher affinity for TβRII. Thus the nuances of differential affinities for TβRI and II, and the disparate levels of expression of essential co-receptors in diverse cell types, determine whether receptor ligation is sufficient to drive signalling. It is also possible that co-receptors directly deliver secondary signals that contribute to target cell activation, a question that we are now investigating.

The affinities of TGM family members for TβRs may thus be calibrated by evolution to depend on interaction with co-receptors such as CD44 that is bound by both TGM1 [15] and TGM4 interactions. Notably, TGM4 interacts with a broader range of co-receptor partners, including CD49d, CD206 and Nrp-1, and while CD44 interactions are governed by D4-5, CD49d interacts with D1-3. This may explain while cell binding by TGM4 is primarily mediated by D4- 5, it is diminished compared to full-length TGM4 in which D1-3 may contribute.

CD49d is an integrin α chain subunit (α4, Itga4), that combines with β integrins; dimerised to β1 integrin (CD29) it comprises the VLA (very late antigen) 4 surface marker that mediates binding to VCAM1 (CD106), expressed on eosinophils, myeloid cells and mesenchymal stem cells, in an interaction controlling leukocyte endothelial adhesion, rolling and extravasation. When combined with β7 integrin, CD49d forms a homing receptor which binds mucosal vascular addressin cell adhesion molecule 1 (MAdCAM) in mucosal tissues. CD49d expression is raised in many stimulated cell types, such as activated eosinophil subsets compared to basal populations [31]. Thus by targeting CD49d, TGM4 can encompass multiple immune cell subsets that are instrumental to protective immunity against infection.

Other integrins facilitate the activation of latent TGF-β, principally αv combined with different β subunits, releasing mature TGF-β from the latency associated peptide within extracellular matrix; however α4 integrin is not implicated in this process [32,33]. As we ascertained that anti-TGF-β antibody did not reduce signal from TGM4, it is unlikely that release of host TGF- β contributes significantly to the effect observed.

Future work will aim not only to provide finer detail on how TGM proteins interface with multiple receptors, at the structural level and in terms of target cell populations, but also to evaluate the in vivo role of these products. Although it is not yet possible to gene target *H. polygyrus* and similar nematode parasites, antibody neutralisation experiments are a feasible approach to test if TGM proteins are essential for helminth survival in the host. Furthermore, TGM1 has proven to be effective in multiple mouse models of inflammation [6–9]; it may be instructive to test TGM4 in similar settings, if selective modulation of the myeloid cell population would offer therapeutic benefit.

## Acknowledgements

SPS, DJS, KC, MPJW, TC, and CC and were supported by the Wellcome Trust through an Investigator Award to RMM (Ref 219530), and the Wellcome Trust core-funded Wellcome Centre for Integrative Parasitology (Ref: 104111); AM, C-HB, CSH and APH by NIH grants R03 AI153915 and F30 AI157069; SL, SZ, CS and GI by the Beatson Institute; and MvD and PtD by Oncode institute base funds. We gratefully acknowledge assistance and expertise from the Flow Core Facility and the Wolfson Research Facility at the University of Glasgow.

## Materials and Methods

### General Materials

Details of enzymes, chemicals, substrates, transfection and labelling reagents and other materials purchased from commercial suppliers is given in **Suppl. Table 3.**

### Expression of TGM1 and TGM4 recombinant proteins

For cellular and *in vivo* applications using live cells, recombinant proteins were expressed in mammalian HEK293 cells. Mammalian codon-optimised genes were synthesized by GeneArt as previously published [14], and subcloned into the mammalian expression vector pSecTag2A using restriction sites *Asc*I and *Not*I, or *Asc*I and *Apa*I. Amplification and cloning of the truncated versions of TGM4 (D1-3 and D4-5) was performed by PCR amplification using proofreading Taq polymerase Phusion Hi as per the manufacturer’s instructions (Invitrogen), full-length codon-optimised TGM4 as template, and the primers shown in **Suppl. Table 4**. For biophysical procedures, individual domains were expressed in *E. coli.* DNA inserts coding for TGM-4 domains D2 and D3 were inserted into a modified pET32a vector between the *NdeI* and *HindIII* sites. Constructs were expressed in BL21 (DE3) cells and cultured at 37°C, to an OD of 0.8, at which point protein expression was induced by adding 0.8 mM isopropyl β-D-1- thiogalactopyranoside (IPTG). Overexpressed TGMs were purified from the conditioned medium by capturing on nickel-loaded HiTrap chelating columns (Cytiva), and after washing until the UV absorbance returned to baseline, the protein was eluted with a 0.0 - 0.5 M imidazole gradient. The fractions with TGM were pooled, concentrated, and further purified on a Superdex 200 16/60 column (Cytiva).

### Production and purification of TGF-β family and CD44 receptor extracellular domains (ECDs)

Biotinylated avi-tagged human TβRI and TβRII were produced as insoluble proteins in *E. coli*, and after reconstitution, refolding, and purification, were enzymatically modified with purified recombinant BirA, as previously described [4]. The human ActRII and BMPRII receptor extracellular domains, preceded by the rat serum albumin signal peptide, a hexahistidine tag and a thrombin cleavage site, were inserted into a pcDNA 3.1+ vector between the *NheI* and *XhoI* sites. The constructs were expressed in expi293 cells (Thermo) and purified from the conditioned medium in an identical manner to that described above for the TGMs. The mouse and human CD44 extracellular domains were expressed in expi293 cells and purified from the conditioned medium as previously described [15].

### Labelling of TGM1/TGM4 for use in flow cytometry

Recombinant TGM1 and TGM4 D1-5, D1-3 and D4-5 proteins were fluorescently labelled with Alexa Fluor™ 594 or 488 Microscale Protein Labeling Kits (Invitrogen™, A30008 or A30006) as described [15]. Briefly, 50 µg (∼1 mg/ml) protein was mixed with Alexa dye and 1 M sodium bicarbonate at 1/10th of the reaction volume concentration, and incubated at room temperature for 15 min. Unlabelled dye was removed from the reaction mixture on a desalting column supplied in the labelling kit. Protein concentrations were calculated using a Nanodrop spectrophotometer (Thermo Scientific). The degree of labelling (DOL, the average number of dye molecules per protein molecule) of the dye-conjugated TGMs was 7.8, 3.6 and 6.8 for TGM-1-AF488, TGM-1-AF594 and TGM-4-AF488 respectively.

### Cell lines

All reagents and cell lines used in this study are listed in **Suppl. Table 5**. HepG2 cells were a kind gift from Dr. Saverio Tardito, CRUK Beatson Institute, Glasgow. All cell lines used in this study were grown and maintained in DMEM supplemented with 10% fetal bovine serum, 2 mM L-glutamine and 100 I.U./ml penicillin-streptomycin in tissue culture dishes or flasks at 37°C, 5% CO2.

### Primary Splenocytes

Spleens recovered from C57BL/6J mice were pressed through a 100 µm strainer, flushed with 20 ml RPMI1640 medium to obtain single cell suspensions. Cells were pelleted at 400 *g,* and resuspended in 1 ml red blood cell lysis buffer (Sigma) for 5 min at room temperature. Cells were washed and resuspended in RPMI1640 medium, and counted using a haemocytometer in diluted trypan blue solution.

### Bone Marrow Derived Macrophages

Bone marrow was recovered from the femurs and tibias of naïve C57BL/6J or BALB/c mice. Briefly, the connective and muscular tissue were removed, bones washed, and the tips of the epiphyses cut using a sterile scissors and forceps. The bone marrow was then flushed out with DMEM using a 25-gauge needle into a petri dish. The flecks of bone marrow were then homogenized using a 19-gauge needle before filtration through a 70 μm filter into a sterile tube. The single cell suspension was then centrifuged at 400 *g* for 5 min at 4°C. Cells were then counted and resuspended at 1 x 10^6^ cells/ml in DMEM supplemented with 30% L929 media containing CSF-1. Cells were then incubated for 6-7 days, with fresh media added on day 3. On the final day, cells were harvested by washing with PBS to remove any potential non-adherent cells, followed by scraping to remove the adherent macrophages. BMDM were then counted, resuspended at the appropriate concentration and plated in 96-well plates for ELISA experiments, or 24-well plates for flow cytometric experiments. Cells were allowed to rest for 1-2 h before stimulation to allow for adherence to the new plate.

### Reporter Bioassays

The TGF-β bioassay with MFB-F11 cells developed by Tesseur et al. [18] was performed as previously described [3]. MFB-F11 cells were tested and found to be mycoplasma-free. Briefly, confluent cells were detached with trypsin, and resuspended in DMEM with 2.5% FCS, 100 U/ml of penicillin, 100 μg/ml of streptomycin and 2 mM L-glutamine at a concentration of 8×10^5^ cells/ml. In 50 μl, 4×10^4^ cells were added to each well of a 96-well round-bottomed plate. Dilutions of purified proteins were then added to each well in a volume of up to 50 μl and incubated for 24 h at 37 °C. Subsequently, 20 μl of supernatant were aspirated from each well, added to an ELISA plate (Nalge Nunc International, USA) with 180 μl of reconstituted Sigma FastTM p-nitrophenyl phosphate substrate and incubated at room temperature in the dark for up to 4 h. Plates were read on at 405 nm on an Emax precision microplate reader (Molecular Devices, USA). All conditions were set up in duplicate and repeated at least twice.

### Western blotting

Cell lysates and pull-down samples were analysed on 4-12% bis-tris SDS-PAGE gels and transferred onto nitrocellulose membrane using iBlot2 (Invitrogen, IB21001). Membranes were treated in 5% non-fat milk blocking solution for 1hr and incubated with primary antibodies listed in **Suppl. Table 6** (1:1000 in 5% BSA containing TBST) overnight at 4°C and washed 3x (5 min) using 1x TBST. Fluorescently conjugated secondary antibodies as listed in **Suppl. Table 7**, diluted 1:10000 in 5% BSA containing TBST were used to detect the protein bands by Odyssey CLx Imaging System (LI-COR Biosciences).

### pSMAD stimulation

Cells were cultured in 6-well tissue culture plates (Corning™) until they reached a confluency of 80-90% in complete growth medium (DMEM, 10% FBS, 1% L-glutamine, 1X penicillin-streptomycin). The growth medium was then replaced with serum-free DMEM, and cells were incubated at 37°C with 5% CO2 for 4 hours. BMDM and splenic T cells isolated from C57BL/6 mice were starved in serum free medium for 2h. To stimulate pSMAD2, TGFβ, TGM1 or TGM4 were added to the cells and incubated at 37°C for 1 hour. The cells were washed with ice-cold PBS and lysed with RIPA buffer (0.05M Tris-HCl, pH 7.4, 0.15M NaCl, 0.25% deoxycholic acid, 1% NP-40, 1mM EDTA) containing 1X Halt protease and phosphatase inhibitors (Invitrogen™). Cell lysates were then cleared by centrifugation at 13000 *g*, 4°C for 5 minutes, and protein concentrations were estimated using the Precision Red reagent.

### Foxp3^+^ Treg induction assay

A single cell suspension was prepared from the spleens of naïve BALB/c or Foxp3-GFP BALB/c transgenic mice [34], with 2 min incubation in red blood cell lysis buffer (Sigma). Cells were then washed and resuspended in DMEM containing HEPES (Gibco), supplemented with 2 mM L-glutamine, 100 U/ml of penicillin and 100 μg/ml of streptomycin (Gibco), 10% heat-inactivated FCS (Gibco), and 50 nM 2-mercaptoethanol (Gibco). Naive CD4^+^ T cells were isolated by magnetic sorting using the mouse naïve CD4^+^ T cell isolation kit on the AutoMACS system (Miltenyi, Germany) as per the manufacturer’s instructions. Cells were cultured at 2×10^5^ per well in flat-bottomed 96-well plates (Corning, USA) with the addition of IL-2 (Miltenyi) at a final concentration of 400 U/ml and pre-coated with 10 µg/ml of anti-CD3 (eBioscience). Cells were cultured at 37 °C in 5% CO2 for at least 72 h before being removed for flow cytometric analysis. For TβRI (ALK5) inhibitor assays, 5 µM SB431542 (Tocris Bioscience, UK) was added, with DMSO added to control wells.

### Surface staining with antibodies and labelled TGM

Cells were prepared for flow cytometric analysis in PBS, incubated with Fixable Viability Dye eFluor™ 506 at a dilution of 1:1000 in the dark for 25 min at 4°C. Thereafter, cells were washed twice in FACS buffer. In some assays, LIVE/DEAD® fixable blue (Life Technologies, USA) was diluted to 1:1000 in PBS; 100 μl were added to each sample of cells, which was then incubated in the dark for 20 min at 4°C and washed twice in FACS buffer (1 x PBS, 0.5% (w/v) BSA, 0.05% sodium azide). Fc receptors were blocked by incubating cells with 1:50 anti-mouse CD16/CD32 (Fc block, Invitrogen) for 10 minutes at 4°C, followed by two washes with FACS buffer. Antibodies used are listed in **Suppl. Table 8**. Separate Foxp3 staining was not required as cells were from Foxp3-GFP transgenic mice. Labelled TGMs were serially diluted in brilliant stain buffer and cells incubated for 20 min at 4°C. Cells were washed twice with FACS buffer and filtered before acquisition on a BD FACSCelesta cytometer. Wherever available, isotype controls and fluorescence minus one (FMO) controls were used. Single-stained compensation beads were used for compensation settings. FACS data was analysed using FlowJo and Prism GraphPad software.

### Surface Plasmon Resonance

All SPR experiments were performed with a BIAcore ×100 system (Cytiva). Neutravidin was coupled to the surface of a CM5 chip (Cytiva) by EDC-NHS activation of the chip, followed by injection of neutravidin (Thermo) over the surface in sodium acetate, pH 4.5 until the RU increased by 6000-15000 RU. Biotinylated Avi-tagged TβRI and biotinylated Avi-tagged TβRII were captured onto the chip surface at a maximum density of 150 RU. All experiments were performed in HBS-EP buffer, 25 mM HEPES, 150 mM NaCl, 0.005% P20 surfactant, pH 7.4, at an injection rate of 100 μL min^-1^. The surface was regenerated in between each injection with a 30 second injection of 1 M guanidine hydrochloride. The experimental sensorgrams were obtained with double referencing with a control cell coated similarly with neutravidin but lacking the captured receptor and 8 blank buffer injections at the beginning of the run before injection of the samples. The data was analyzed by fitting the results to a 1:1 kinetic model using the SPR analysis software Scrubber (BioLogic Software).

### Isothermal Calorimetry

All ITC experiments were performed with a Microcal PEAQ-ITC system (Malvern Instruments). All experiments were performed in 25 mM Na2HPO4, 50 mM NaCl, pH 6.0 at 35 °C, with 15 2.5 μL injections with a duration of 5 s, a spacing of 150 s, and a reference power of 10. All samples were dialyzed against the same ITC buffer before loading into the system. Data was globally fit using the programs NITPIC [35], SEDPHAT [36,37], and GUSSI [38].

### Nuclear Magnetic Resonance

NMR samples were prepared in 25 mM Na2HPO4, 50 mM NaCl, pH 6.0. Each sample was run in a 5 mm susceptibility-matched NMR microtube for data collection. NMR data was collected with Bruker 600, 700, or 800 MHz spectrometer containing a 5 mm ^1^H (^13^C, ^15^N) z-gradient “TCI” cryogenically cooled probe at a temperature of 37°C. ^1^H-^15^N HSQC spectra were acquired as described, with water flipback [39] and WATERGATE suppression pulses [40]. NMR spectra were processed, analyzed, and visualized using NMRPipe [41] and NMRFAM-SPARKY [42].

### Cell Transfection

Extrachromosomal expression plasmids were transfected into MFB-F11 cells with Lipofectamine-2000, RAW264.7 cells with Lipofectamine LTx and HepG2 with Lipofectamine 3000 according to manufacturer’s instruction. In 6 well plates, 3×10^5^ cells were allowed to adhere overnight. 6 μl lipofectamine reagent and 2.5 μg plasmid DNA diluted and mixed in 200μl serum free DMEM in two separate tubes. Later, diluted plasmid DNA and lipofectamine were pooled together and mixed well and incubated at room temperature (10min). This mixture was added to the cells and mixed by swirling the plate. Transfection to cells to be used for GFP-TRAP pull down were performed in 15 cms tissue culture plates.

### Molecular Cloning and CRISPR-Cas9 Knockout

To engineer pSecTag2A-eGFP plasmid, eGFP was PCR amplified from pEGFP-N1 using primers described in **Suppl. Table 9** and inserted in pSecTag2A plasmid using *PstI* and *XhoI*. TGM-1 D1-5, TGM4 D1-5, D1-3 and D4-5 coding sequences were PCR amplified using primers mentioned in **Suppl. Table 9** and were inserted using *AscI* and *NotI* into the pSectAg2A-eGFP plasmid to generate pSecTag2A-TGM-eGFP constructs.

For deletion of CD44, CD49d and NRP1 expression in Raw264.7 macrophage or MFB-F11 fibroblast cell lines, a CRISPR strategy was used [43], with guides mentioned in **Suppl. Table 9** cloned in pSpCas9(BB)-2A-GFP (a kind gift from Dr. Jamie Whitelaw, CRUK Beatson institute, Glasgow); Addgene plasmid #48138). 2.5 mg empty and CD44 guide RNA containing pSpCas9(BB)-2A-GFP plasmids were transfected in MFB-F11 or RAW264.7 cells. 24 h post transfection GFP positive single cells were sorted by FACS in 96-well plates. Single cell clones were screened by Western blotting after approximately 2 weeks.

### GFP-TRAP pull down

For expression of eGFP fusion proteins, MFB-F11, RAW and HepG2 cells were grown to 70% confluence in 150 mm Petri dishes. Approximately 50 µg of pSecTag2A-eGFP, or the pSecTag2A-eGFP plasmid containing in-frame fusions with TGM1 and TGM4 full-length (FL), D1-3 or D4-5 were transfected into MFB-F11 or RAW or HEPG2. 40 hr post transfection cells were lysed with cell lysis buffer (10 mM Tris-HCl pH 7.5, 150 mM NaCl, 0.5 mM EDTA and 0.5% NP-40) supplemented with 1X Halt protease and phosphatase inhibitor cocktails (Invitrogen). Cell lysates were cleared by centrifugation (13000 *g*, 10 min). 2 mg of cell lysates were incubated with 25 ml of GFP-TRAP beads (Chromotek, GTA-20) for 1 hr at 4°C on rotation. Beads were washed 4X with cell-lysis buffer (5 min each wash on rotation). To elute the proteins from the beads, 50 μL 2 x NuPAGE LDS sample buffer with 25mM DTT was added and boiled (100°C, 5 min). Protein samples were analyzed on 4-12% Bis-tris acrylamide gels followed by Western blotting.

### Streptavidin pull down

Ten µg of TGM1 and TGM4 D1-5, D1-3 and D4-5 were biotinylated and purified as described [15]. For the pull down, MFB-F11, RAW264.7 and CD44 knock out cells were grown at 80- 90% confluency in 15 cm tissue culture dishes, washed 3x with ice-cold PBS and incubated with ∼3.5 µg of biotinylated TGMs for 3 hr on ice. Cells were washed 3x with ice-cold PBS and lysed with Cell Lysis Buffer (100 mM NaCl, 25 mM Tris, pH7.5, 5 mM MgCl2 and 0.5% NP40) supplemented with 1x Halt protease inhibitor (Thermo Scientific, 1861279) and phosphatase inhibitor (Thermo Scientific, 78427) cocktails. Cell lysates were cleared by centrifugation (13000 *g*, 10mins). Two µg of cell lysates were incubated with 30 µl of Neutravidin agarose beads (Thermo Scientific, 29201) for 1 hr at 4°C. Beads were washed with lysis buffer 4x (5 min each). For mass-spectromertry beads were stored in 100 mM ammonium bicarbonate at -20°C. For western blotting, 50 µl LDS sample buffer (Invitrogen, NP0007) containing 25 mM dithiothreitol (DTT) was added to beads and heated for 5 min at 100°C.

### Liquid Chromatography and Mass-spectrometry (LC-MS)

Neutravidin agarose beads were resuspended in a 2 M Urea and 100 mM ammonium bicarbonate buffer and stored at -20oC. Three biological replicates for each condition were digested with Lys-C (Alpha Laboratories) and trypsin (Promega) “on beads” as previously described [44]. Peptides resulting from all trypsin digestions were separated by nanoscale C18 reverse-phase liquid chromatography using an EASY-nLC II 1200 (Thermo Scientific) coupled to an Orbitrap Q-Exactive HF mass spectrometer (Thermo Scientific). Elution was carried out at a flow rate of 300 nl/min using a binary gradient, into a 20 cm fused silica emitter (New Objective) packed in-house with ReproSil-Pur C18-AQ, 1.9 μm resin (Dr Maisch GmbH), for a total run-time duration of 125 minutes. Packed emitter was kept at 35 °C by means of a column oven (Sonation) integrated into the nanoelectrospray ion source (Thermo Scientific). Eluting peptides were electrosprayed into the mass spectrometer using a nanoelectrospray ion source. An Active Background Ion Reduction Device (ESI Source Solutions) was used to decrease air contaminants signal level. The Xcalibur 4.2 software (Thermo Scientific) was used for data acquisition. A full scan was acquired at a resolution of 120000 at 200 m/z, over mass range of 350-1400 m/z. HCD fragmentation was triggered for the top 15 most intense ions detected in the full scan. Ions were isolated for fragmentation with a target of 1E5 ions, for a maximum of 125 ms, at a resolution of 15,000 at 200 m/z. Ions that have already been selected for MS2 were dynamically excluded for 20 sec.

### MS Data Analysis

The MS Raw data were processed with MaxQuant software [45] version 1.6.14.0 and searched with Andromeda search engine [46], querying SwissProt [47] *Mus musculus* (25198 entries). First and main searches were performed with precursor mass tolerances of 20 ppm and 4.5 ppm, respectively, and MS/MS tolerance of 20 ppm. The minimum peptide length was set to six amino acids and specificity for trypsin cleavage was required. Cysteine carbamidomethylation was set as fixed modification, whereas Methionine oxidation, Phosphorylation on Serine-Threonine-Tyrosine, and N-terminal acetylation were specified as variable modifications. The peptide, protein, and site false discovery rate (FDR) was set to 1%. All MaxQuant outputs were analysed with Perseus software version 1.6.13.0 [48].

Protein abundance was measured using label-free quantification (LFQ) intensities reported in the ProteinGroups.txt file. Only proteins quantified in all replicates in at least one group, were measured according to the label-free quantification algorithm available in MaxQuant [49]. Missing values were imputed separately for each column, and significantly enriched proteins were selected using a permutation-based Student’s t-test with FDR set at 5%.

### Quantification, statistical analysis and software

Western blotting was quantified using ImageJ (FIJJI). FACS data were analysed with FlowJo. Unpaired student’s t-Test or ANOVA was performed using Graph Pad Prism. A full list of software used in this study is presented in **Suppl. Table 10.**

### DATA availability

All raw data and reagents, including plasmids generated by this study (**Suppl. Table 11**) can be requested from the authors.

## Supplementary Material

### Supplementary Figures

**Supplementary Figure 1.**
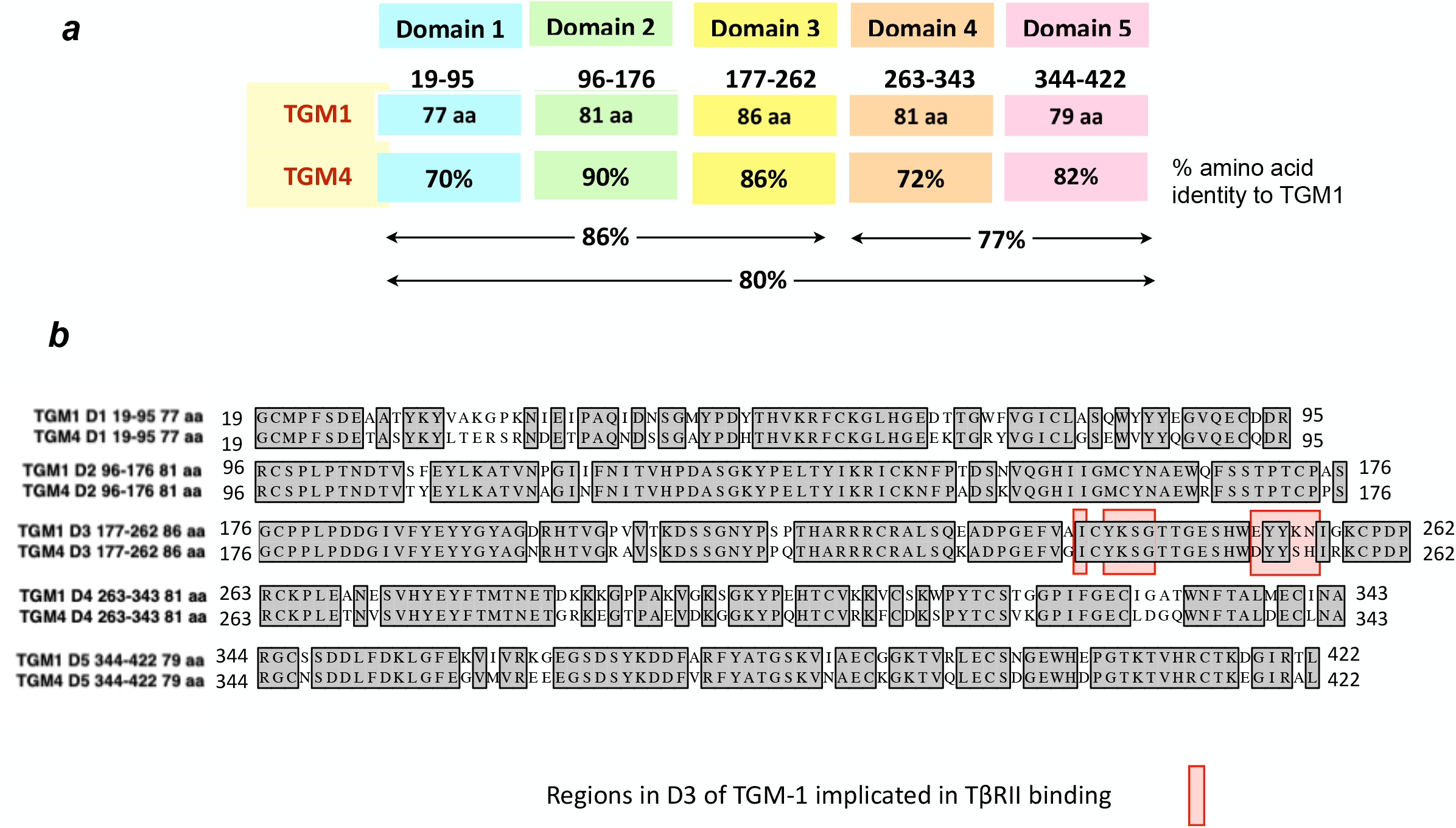
Schematic Structure and Similarity of TGM1 and TGM4 domains. ***a.*** Schematic of domain organisation; figures denote amino acid identity for each domain to TGM1, and amino acid identity for D1-3, D4-5 and full-length TGM4. ***b.*** Amino acid aligments for each domain of TGM1 and TGM4; identical domains are shaded. Red background denotes regions of TGM1 identified as contacting TβRII.

**Supplementary Figure 2.**
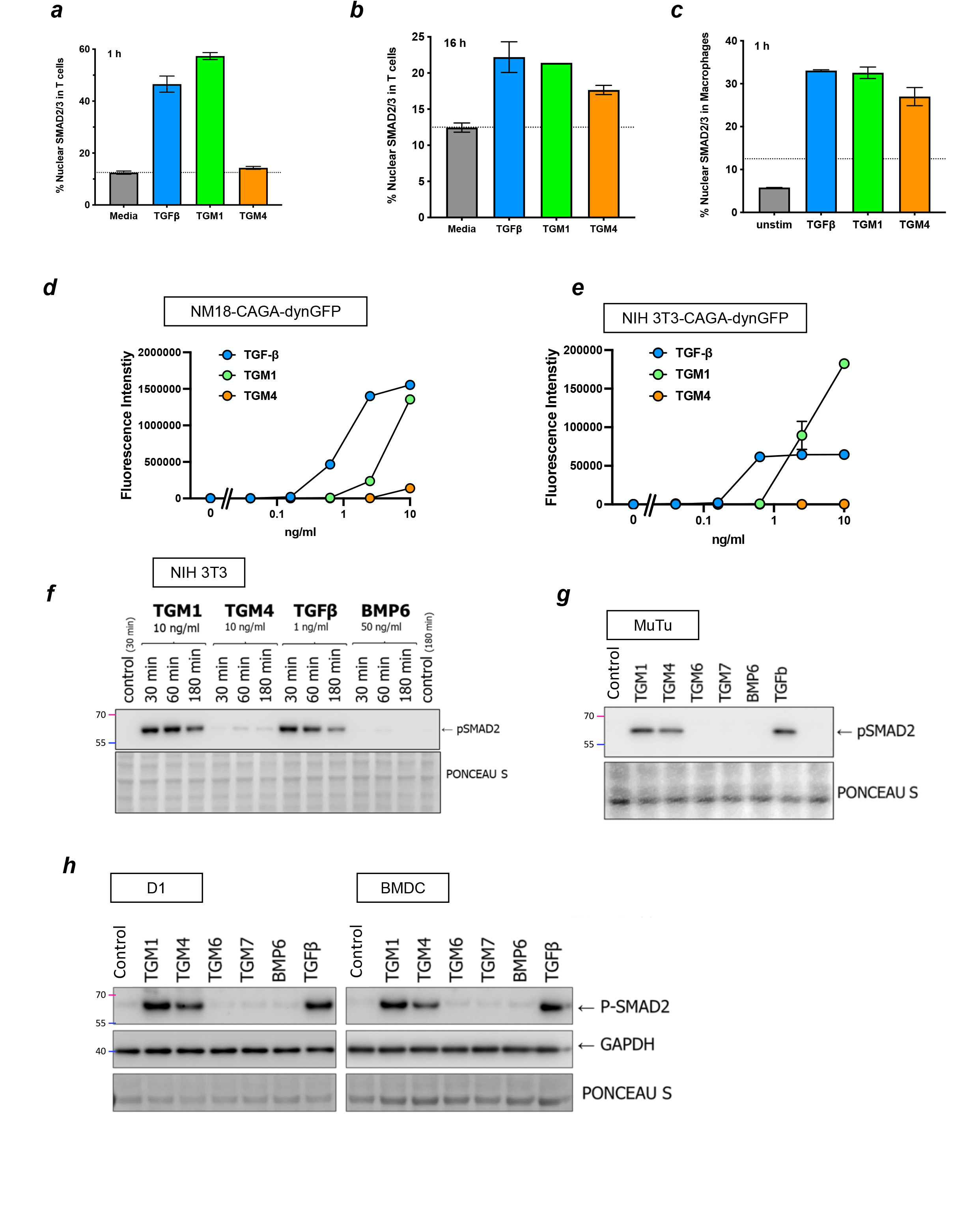
Responses of Different Cell Types to TGM4. ***a-c.*** SMAD2/3 nuclear localization by imaging flow cytometry in T cells at 1 hr (***a***) and 16 hr (***b***) post-stimulation, and in macrophages at 1 hr (***c***), stimulated with TGFβ, TGM1 or TGM4, evaluated by ImageStream. ***d,e.*** NM18 mouse mammary gland epithelial cell (***d)*** and NIH 3T3 mouse embryonic fibroblast (***e***) lines transfected with the CAGA-dynGFP reporter plasmid, and stimulated with TGFβ, TGM1 or TGM4, assayed by fluorescent intensity at 24 hr. f. NIH 3T3 cells analyzed for p-SMAD induction by Western blot by the indicated concentrations of TGFβ, TGM1 or TGM4, for 30, 60 or 180 mins. ***g,h***. pSMAD induction in MuTu mouse splenic dendritic cells. Cells were stimulated for one hour with 10 ng/ml of each TGM protein, 50 ng/ml BMP6 and 5 ng/ml TGF-β. ***h***. pSMAD induction in the D1 mouse dendritic cell line (left) and Bone Marrow-Derived DCs, differentiated in vitro with GM-CSF(right). Cells were stimulated for one hour with 10 ng/ml of each TGM protein, 50 ng/ml BMP6 and 5 ng/ml TGF-β.

**Supplementary Figure 3.**
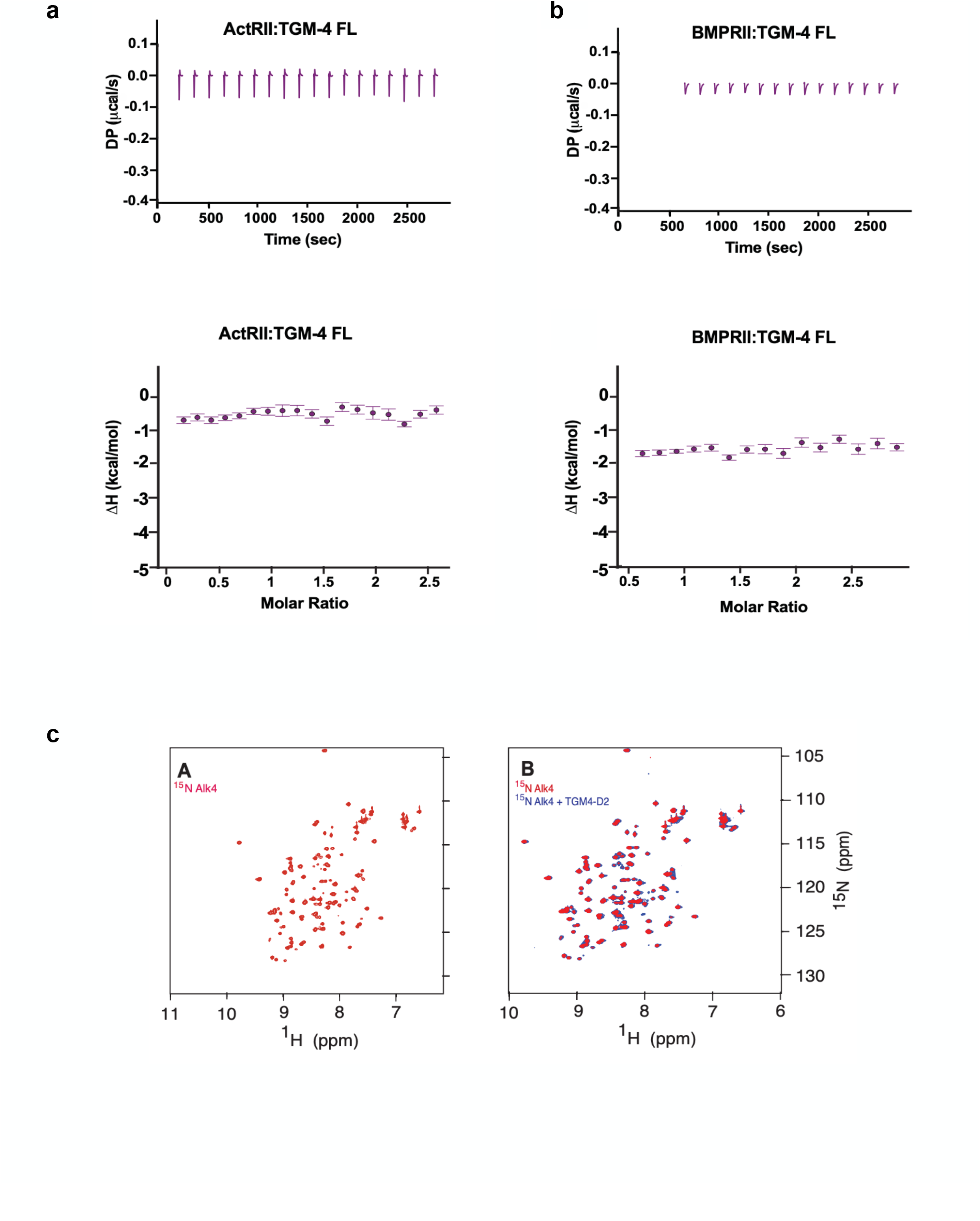
Testing binding of TGM4 to other TGF-β family receptors. ***a.*** ITC analysis of FL TGM4 interactions with ActRII; upper panel present the raw thermograms for the injection of ActRII into a cell containing TGM-4 FL; lower panel presents the integrated heats for these data. ***b.*** As ***a***, for interactions of TGM4 with BMPRII. ***c.*** NMR analysis of TGM4 D2 interaction with ALK4 Type I receptor. ^1^H-^15^N spectrum of ^15^N Alk4 alone (left, red) and overlaid onto the ^1^H-^15^N spectrum of ^15^N Alk4 bound to 1.2 molar equivalents of unlabeled TGM4 D2 (right, blue). The spectra were recorded in 25 mM sodium phosphate, 50 mM sodium chloride, 5% ^2^H2O, pH 6.0 at a sample temperature of 37°C.

**Supplementary Figure 4.**
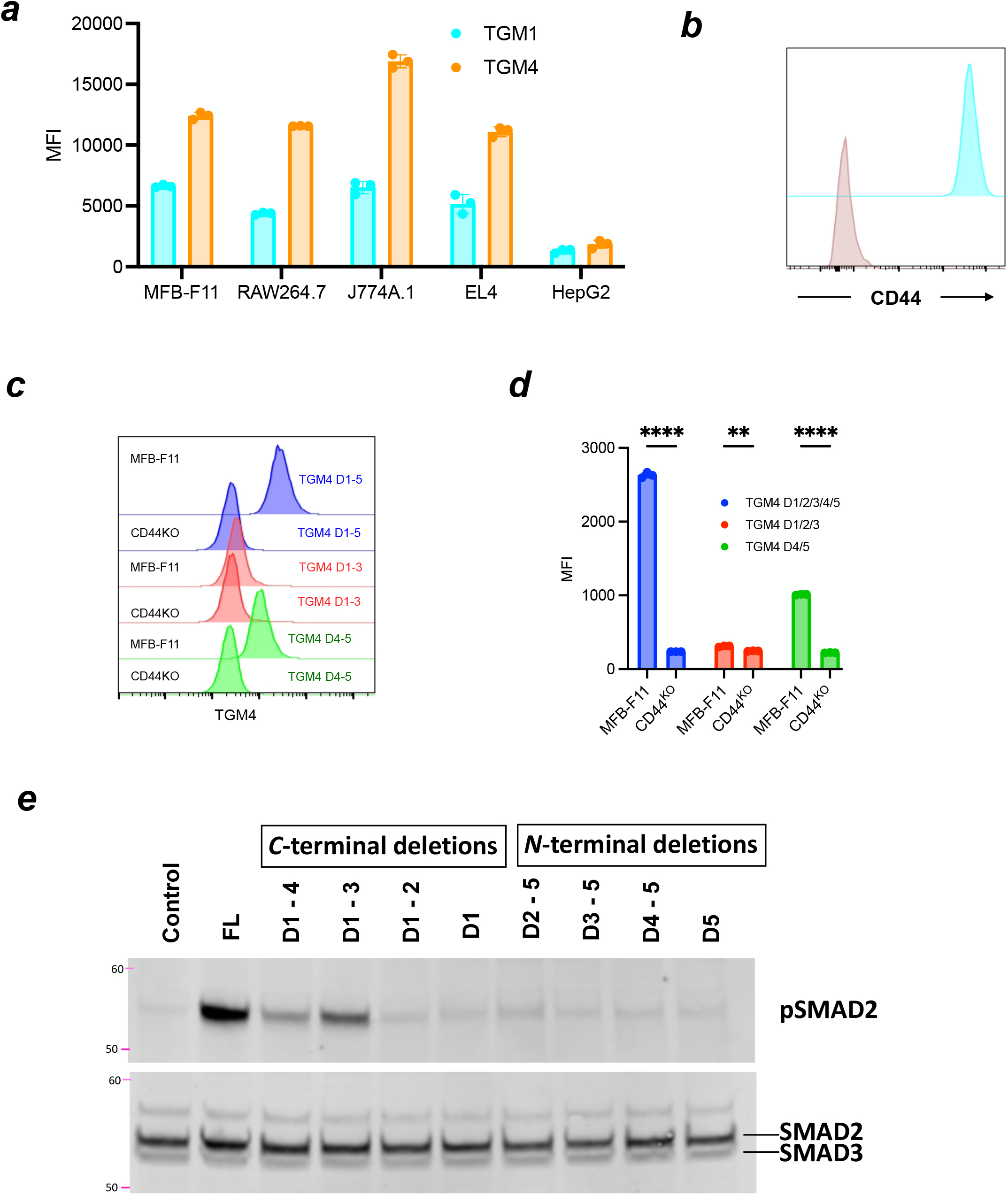
Domain and Co-Receptor Interactions of TGM4. ***a.*** Binding of AF-594-labelled TGM1 and TGM4 constructs to the indicated cell lines, measured by Mean Fluorescence Intensity on a flow cytometer. ***b.*** Flow cytometric analysis of CD44 binding to control RAW264.7 cells (cyan) and Cd44- deleted RAW264.7 cells (tan). ***c,d.*** Flow cytometric analysis of TGM4 binding to MFB-F11 wild-type and CD44-deficient cells, probed with full length TGM4 D1-5, and truncated constructs D1-3 and D4-5. Example histograms (***c***) and results from 3 replicate experiments (***d***) are shown. **, p<0.01; ****, p<0.0001. ***e.*** pSMAD induction by truncated TGM4 constructs, with C-terminal deletions and N-terminal deletions, assessed by Western blot. FL, Full-length.

**Supplementary Figure 5.**
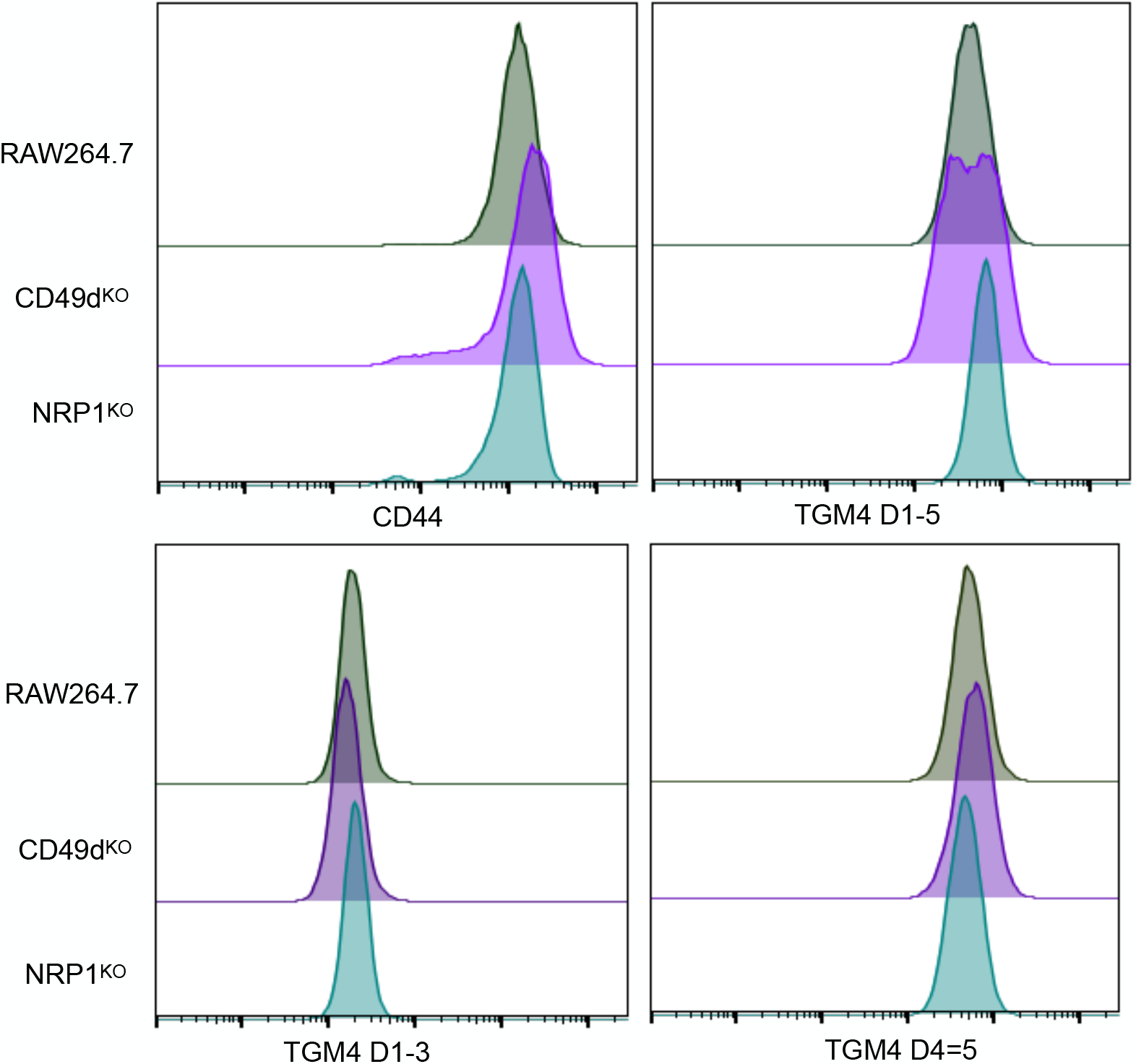
TGM4 Binding to Cells Lacking CD49d or NRP1. RAW264.7 control cells, and sublines in which expression of CD49d or NRP1 has been genetically deleted, were probed by flow cytometry for bidning to anti-CD44, TGM4 D-15, D1-3 or D4-5 as indicated.

**Supplementary Figure 6.**
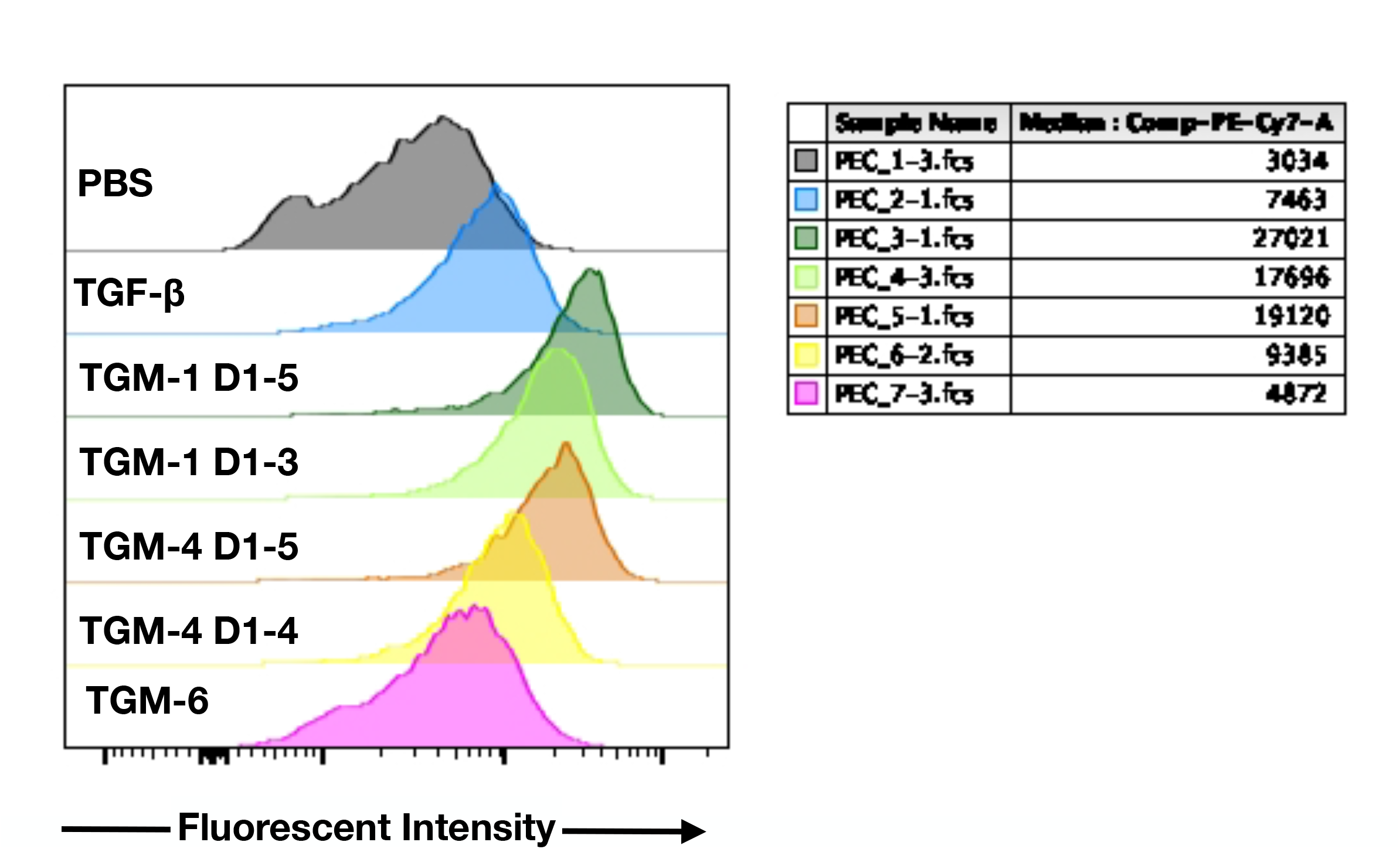
Induction of Arginase-1 by Full-Length TGM1 and TGM4. Flow cytometric histograms of binding by the indicated ligands, from one of the replicates shown in **Figure 7i**.

---

**Suppl. Table 1.** TGM-1 and TGM-4 binding to TβRI and TβRII as assessed by SPR.

| Surface | Analyte (Conc.) | Fitted Parameters<sup>1</sup> |  |  |  |
| --- | --- | --- | --- | --- | --- |
| | | $k_{on}$ ($M^{-1} cm^{-1}$) | $k_{off}$ ($s^{-1}$) | $K_D$ | $R_{max}$ (RU) |
| TβRI | TGM1 (31.25 -1000 nM) | $(6.4 \pm 0.6) \times 10^4$ | $(4.4 \pm 0.1) \times 10^{-3}$ | $70 \pm 6$ nM | $232 \pm 2$ |
| TβRI | TGM4 (62.4 - 250 nM) | $(2.2 \pm 0.1) \times 10^5$ | $(7.3 \pm 0.1) \times 10^{-4}$ | $3.2 \pm 0.1$ nM | $222 \pm 8$ |
| TβRII | TGM4 (12.5 – 100 μM) | $(7.2 \pm 0.2) \times 10^3$ | $(0.84 \pm 0.03) \times 10^0$ | $116 \pm 2$ μM | $141 \pm 2$ |
| <sup>1</sup>Fitted parameters were derived from kinetic analysis of a single injection series |  |  |  |  |  |

**Suppl. Table 2.** TGM-4:CD44 binding as assessed by ITC.

| Cell | TGM4-D4/5 | TGM4-D4/5 | TGM1-D4/5 | TGM1-D4/5 |
| --- | --- | --- | --- | --- |
| Syringe | mCD44 | hCD44 | mCD44 | hCD44 |
| Cell Conc. (μM ) | 5.5 | 5 | 12 | 8 |
| Syringe Conc. (μM ) | 60 | 75 | 90 | 100 |
| Temperature (°C) | 35 | 35 | 35 | 35 |
| $K_D$ (nM) | 20 (7, 41)<sup>ab</sup> | 56 (28, 99)<sup>ab</sup> | 62 (28, 117)<sup>ab</sup> | 140 (86, 221)<sup>ab</sup> |
| $\Delta H$ (kcal mol<sup>-1</sup>) | -21.7 (-23.0, -20.5)<sup>ab</sup> | -16.6 (-17.9, -15.5)<sup>ab</sup> | -19.4 (-20.5, -18.4)<sup>ab</sup> | -9.6 (-10.2, -9.0)<sup>ab</sup> |
| $\Delta G$ (kcal mol<sup>-1</sup>) | -10.9<sup>ab</sup> | -10.2<sup>ab</sup> | -10.2<sup>ab</sup> | -9.7<sup>ab</sup> |
| $-T\Delta S$ (kcal mol<sup>-1</sup>) | 10.8<sup>ab</sup> | 6.4<sup>ab</sup> | 9.2<sup>ab</sup> | -0.1<sup>ab</sup> |
| Stoichiometry (n) | 1.2<sup>d</sup> | 1.5<sup>d</sup> | 1.4<sup>d</sup> | 1.2<sup>d</sup> |

<sup>a</sup>Not determined due to weak signal<sup>b</sup>Uncertainty reported as 68.3% confidence interval<sup>c</sup>Global fit of two replicates<sup>d</sup>Number of sites determined by incompetent fraction value on sedphat; set to '1' for $K_D$ analysis

**Suppl. Table 3.** List of commercially supplied reagents used.

| Sl. no | Name of consumables | Supplier | Cat. no. |
| --- | --- | --- | --- |
| 1. | GFP-TRAP beads | Chromotek | GTA-20 |
| 2. | Halt™ Phosphatase Inhibitor Cocktail | Thermo Scientific™ | 78427 |
| 3. | Halt™ Protease Inhibitor Cocktail (100X) | Thermo Scientific™ | 78438 |
| 4. | EZ-Link™ Sulfo-NHS-LC-Biotin, No-Weigh™ Format | Thermo Scientific™ | A39257 |
| 5. | Pierce™ NeutrAvidin™ Agarose | Thermo Scientific™ | 29201 |
| 6. | Cell Lysis Buffer (10X) | CST | 9803 |
| 7. | Lipofectamine™ 2000 Transfection Reagent | Invitrogen™ | 11668500 |
| 8. | Lipofectamine™ 3000 Transfection Reagent | Invitrogen™ | L3000150 |
| 9. | Lipofectamine™ LTX Reagent | Invitrogen™ | 15338500 |
| 10. | Alexa Fluor™ 488 Microscale Protein Labeling Kit | Invitrogen™ | A30006 |
| 11. | Alexa Fluor™ 594 Microscale Protein Labeling Kit | Invitrogen™ | A30008 |
| 12. | DTT | Melford | 3483-12-3 |
| 13. | Precision Red Advanced Protein Assay: 1x stock | Cytoskeleton | ADV02 |
| 14. | PD MiniTrap desalting columns with Sephadex G-25 resin | Cytiva | 28918007 |
| 15. | One Shot™ Stbl3™ Chemically Competent *E. coli* | Invitrogen™ | C737303 |
| 16. | VeriFi™ Hot Start Mix | PCR Biosystems | PB10.46-01 |
| 17. | T4 Ligase | NEB |  |
| 18. | T4 Polynucleotide Kinase | NEB | M0201S |
| 19. | Adenosine 5'-Triphosphate (ATP) | NEB | P0756S |
| 20. | PstI | NEB | R0140S |
| 21. | XhoI | NEB | R0146S |
| 22. | Ascl | NEB | R0558S |
| 23. | NotI | NEB | R0189S |
| 24. | FastDigest Bpil | Thermo Scientific™ | FD1014 |
| 25. | Tango Buffer (10X) | Thermo Scientific™ | BY5 |
| 26. | Novex™ Sharp Pre-stained Protein Standard | Invitrogen™ | LC5800 |
| 27. | GeneRuler 1 kb DNA Ladder, ready-to-use | Thermo Scientific™ | SM0313 |
| 28. | Pierce™ ECL Plus Western Blotting Substrate | Thermo Scientific™ | 32132 |
| 29. | CM-5 SPR sensor chip | Cytiva | BR100012 |
| 30. | Neutravidin | Thermo Scientific™ | 31000 |

**Suppl. Table 4.**
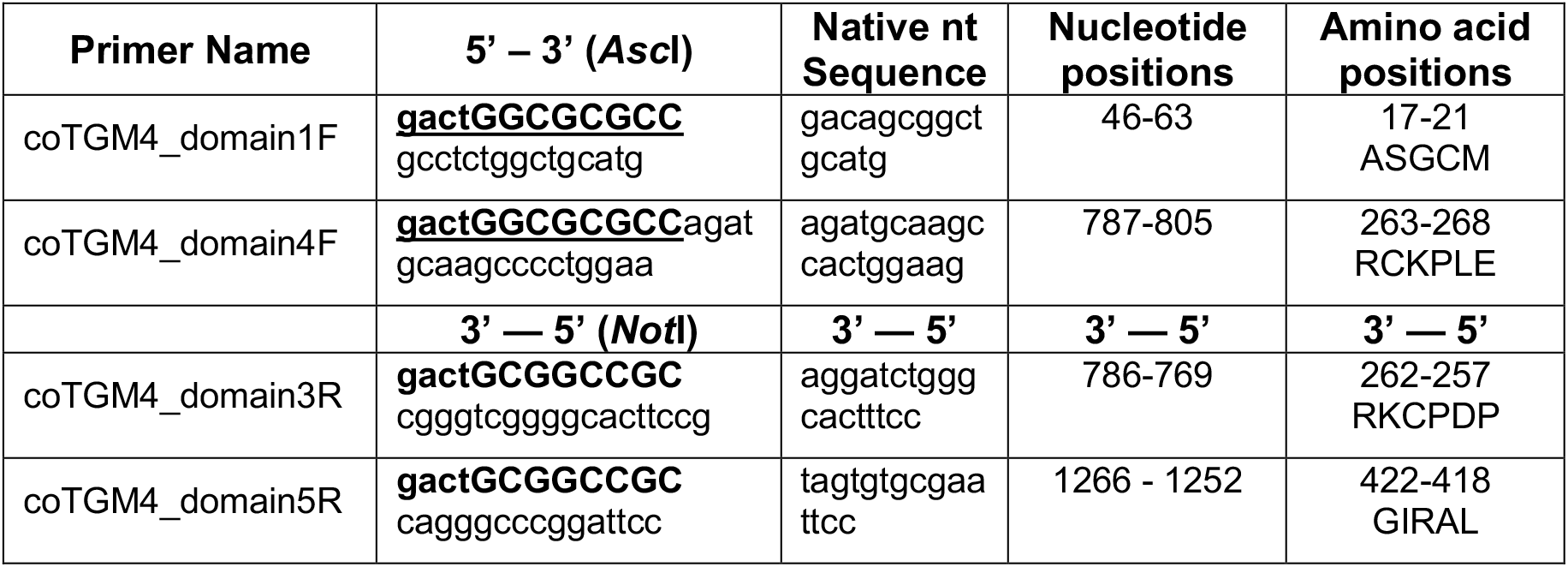
Primers used for PCR amplification of truncated TGM4 proteins D1-3 and D4-5.

For each primer, the gact cap is shown in lower case bold; the restriction sites (GG^CGCGCC for *Asc*I and GC^GGCCGC for *Not*I) in upper case bold, and the sequence corresponding to the domain to be amplified in plain lower case. The third column gives the native *H. polygyrus* TGM4 nucleotide sequence (Genbank Accession number MG429739) for the same segment. The right-hand columns denote the nucleotide positions in the full-length TGM4 open reading frame to which these sequences correspond, and finally the amino acid positions in the full-length TGM4 protein. Amino acid positions are given 5’ to 3’ for both 5’-3’ and 3’-5’ primers

**Suppl. Table 5.** List of cell lines used in this study.

| Sl. no. | Name of the cell line | Source/ATCC |
| --- | --- | --- |
| 1. | MFB-F11 | [18] |
| 2. | RAW264.7 | TIB71 |
| 3. | J774A.1 | TIB67 |
| 4. | EL4 | TIB39 |
| 5. | HepG2 | Kind Gift from CRUK Beatson institute, HB8065 |
| 6. | CD44<sup>KO</sup> in MFB-F11 | In this study |
| 7. | CD44<sup>KO</sup> in RAW264.7 | In this study |
| 8. | CD49d<sup>KO</sup> in RAW264.7 | In this study |
| 9. | NRP1<sup>KO</sup> in RAW264.7 | In this study |

**Suppl. Table 6.** List of primary antibodies used.

| Sl.no. | Antigen | Name | Supplier | Cat. no |
| --- | --- | --- | --- | --- |
| 1. | TGF-β receptor I | Recombinant Anti-TGF-β Receptor I antibody | Abcam | ab235578 |
| 2. | TGF-β receptor II | Recombinant Anti-TGF-β Receptor II antibody | Abcam | ab25936 |
| 3. | CD44 | Recombinant Anti-CD44 antibody |  | ab189524 |
| 4. | CD44 | Recombinant Anti-CD44 antibody |  | ab243894 |
| 5. | CD49d | CD49d (Integrin $\alpha$ 4) Recombinant Rabbit Monoclonal Antibody | Thermo fischer scientific | MA5-35435 |
| 6. | CD206(MRC1) | CD206/MRC1 (E6T5J) XP® Rabbit mAb | CST | #24595 |
| 7. | GFP | ChromoTek GFP Monoclonal antibody | Chromotek | 3H9 |
| 8. | CD72 | Anti-CD72 antibody | Abcam | Ab201079 |
| 9. | SMAD2/3 | SMAD2/3 Antibody #3102 | CST | #3102, #5678 |
| 10. | pSMAD2 | Phospho-SMAD2 (Ser465/467)/SMAD3 (Ser423/425) (D27F4) Rabbit mAb | CST | #8828 |
| 11. | LILRB3 | LILRB3 Polyclonal Antibody |  | PA590933 |
| 12. | NRP1 | Neuropilin 1 | Abcam | ab81321 |
| 13. | His | Anti-His HRP Conjugate Kits | Qiagen | 1014992 |
| 14. | α-Tubulin | Anti-α-Tubulin antibody (DM1A) | Abcam | ab7291 |

**Suppl. Table 7.**
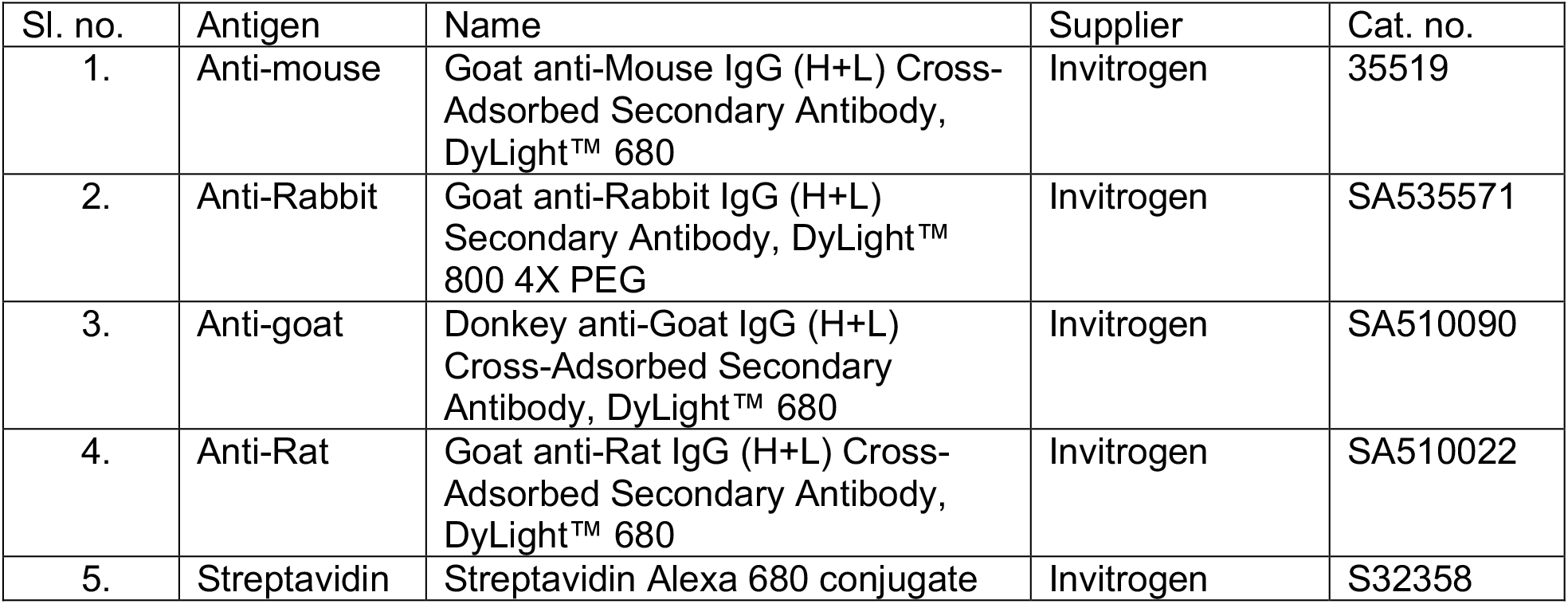
List of secondary antibodies used.

| Sl. no. | Antigen | Name | Supplier | Cat. no. |
| --- | --- | --- | --- | --- |
| 1. | Anti-mouse | Goat anti-Mouse IgG (H+L) Cross-Adsorbed Secondary Antibody, DyLight™ 680 | Invitrogen | 35519 |
| 2. | Anti-Rabbit | Goat anti-Rabbit IgG (H+L) Secondary Antibody, DyLight™ 800 4X PEG | Invitrogen | SA535571 |
| 3. | Anti-goat | Donkey anti-Goat IgG (H+L) Cross-Adsorbed Secondary Antibody, DyLight™ 680 | Invitrogen | SA510090 |
| 4. | Anti-Rat | Goat anti-Rat IgG (H+L) Cross-Adsorbed Secondary Antibody, DyLight™ 680 | Invitrogen | SA510022 |
| 5. | Streptavidin | Streptavidin Alexa 680 conjugate | Invitrogen | S32358 |

**Suppl. Table 8.** List of antibodies and reagents used in Flow-cytometry.

| Sl. no. | Antigen | Name | Supplier | Cat. no. |
| --- | --- | --- | --- | --- |
| 1. | CD44 | FITC anti-mouse/human CD44 Antibody | Biolegend | 103006 |
| 2. | CD44 | PE anti-mouse/human CD44 Antibody | Biolegend | 103008 |
| 3. | CD4 | FITC anti-mouse CD4 Antibody | Biolegend | 100406 |
| 4. | CD45 | PE/Dazzle™ 594 anti-mouse CD45 Antibody | Biolegend | 103146 |
| 5. | Rat IgG | IgG from rat serum | Sigma | I4131 |
| 6. | Compensation beads | UltraComp eBeads™ Compensation Beads | Invitrogen™ | 01-2222-42 |

**Suppl. Table 9.**
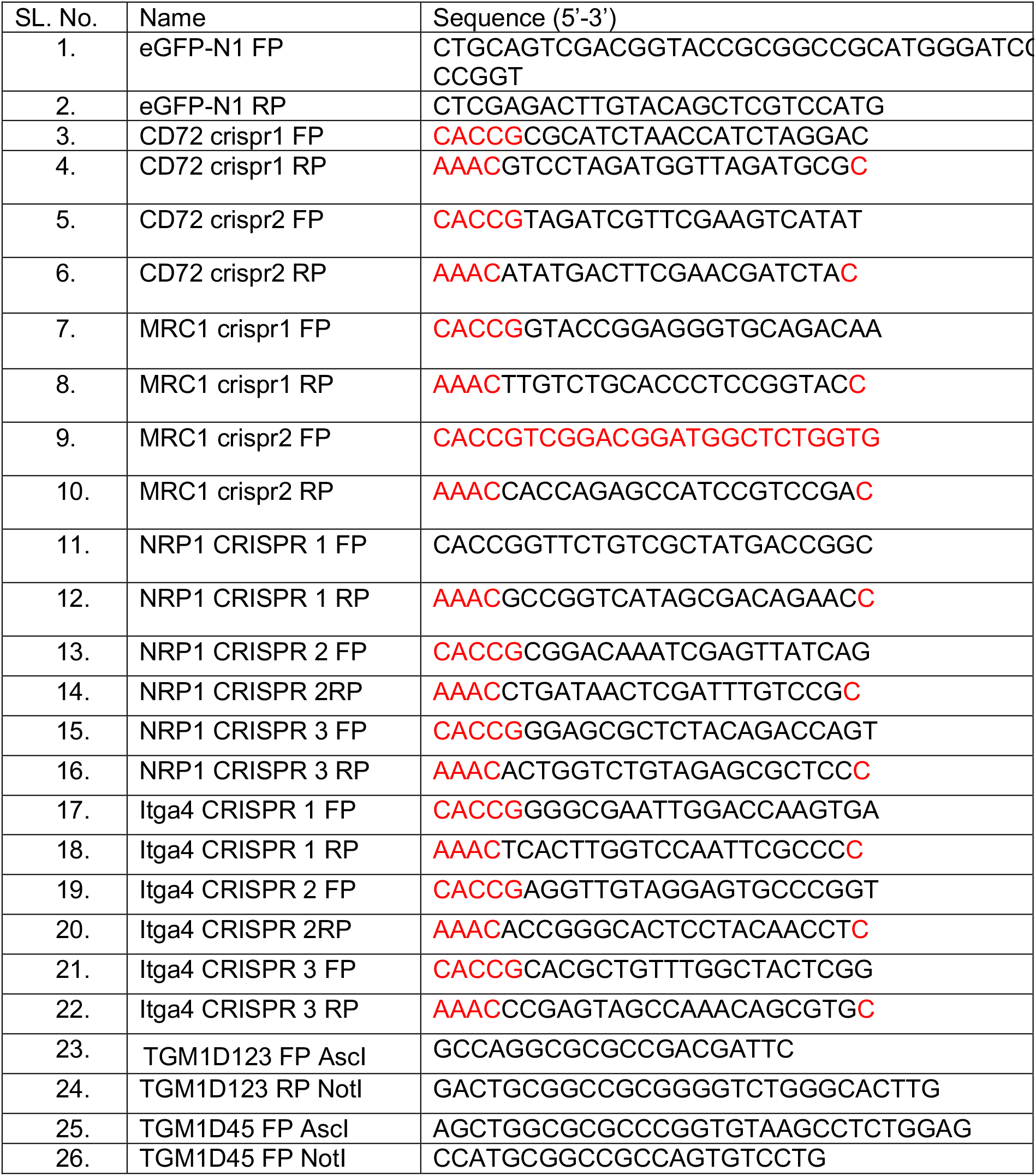
List of oligonucleotides used.

**Suppl. Table 10:**
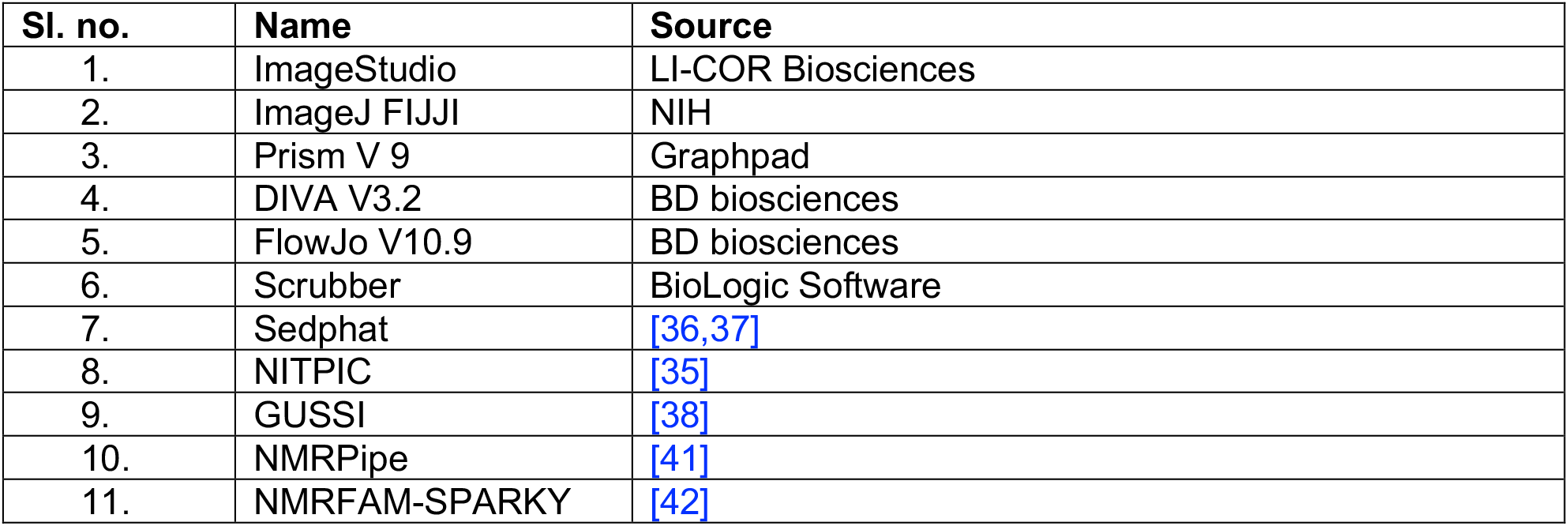
List of software used in this study.

| Sl. no. | Name | Source |
| --- | --- | --- |
| 1. | ImageStudio | LI-COR Biosciences |
| 2. | ImageJ FIJJI | NIH |
| 3. | Prism V 9 | Graphpad |
| 4. | DIVA V3.2 | BD biosciences |
| 5. | FlowJo V10.9 | BD biosciences |
| 6. | Scrubber | BioLogic Software |
| 7. | Sedphat | [36,37] |
| 8. | NITPIC | [35] |
| 9. | GUSSI | [38] |
| 10. | NMRPipe | [41] |
| 11. | NMRFAM-SPARKY | [42] |

**Suppl. Table 11:** List of plasmids used and generated in this study.

| Sl. no. | Name | Description | Source |
| --- | --- | --- | --- |
| 1. | pSpCas9(BB)-2A-GFP | CRISPR vector | PX458; Addgene |
| 2. | pSec-Tag-2A | Modular expression vector | V90020; Thermo Scientific™ |
| 3. | pSP503 | GFP expression vector | In this study |
| 4. | pSP504 | Vector for C-terminal GFP tagging | In this study |
| 5. | pSP505 | TGM1-eGFP expression vector | In this study |
| 6. | pSP506 | TGM4-eGFP expression vector | In this study |
| 7. | pSP508 | TGM1D123-eGFP expression vector | In this study |
| 8. | pSP509 | TGM1D45-eGFP expression vector | In this study |
| 9. | pSP510 | TGM4D123-eGFP expression vector | In this study |
| 10. | pSP511 | TGM4D45-eGFP expression vector | In this study |
| 11. | Neuropilin 1 CRISPR vectors | Neuropilin 1 CRISPR vectors | In this study |
| 12. | CD49d CRISPR vectors | CD49d CRISPR vectors | In this study |
| 13. | CD72 CRISPR vectors | CD72 CRISPR vectors | In this study |
| 14. | CD206 CRISPR vectors | CD206 CRISPR vectors | In this study |

